# Colistin kills bacteria by targeting lipopolysaccharide in the cytoplasmic membrane

**DOI:** 10.1101/479618

**Authors:** Akshay Sabnis, Katheryn L. H. Hagart, Anna Klöckner, Michele Becce, Lindsay E. Evans, R. Christopher D. Furniss, Despoina A. I. Mavridou, Ronan Murphy, Molly M. Stevens, Jane C. Davies, Gerald J. Larrouy-Maumus, Thomas B. Clarke, Andrew M. Edwards

## Abstract

Colistin is an antibiotic of last resort, but has poor efficacy and resistance is a growing problem. Whilst it is well established that colistin disrupts the bacterial outer membrane by selectively targeting lipopolysaccharide (LPS), it was unclear how this led to bacterial killing. We discovered that MCR-1 mediated colistin resistance is due to modified LPS at the cytoplasmic rather than outer membrane. In doing so, we also demonstrated that colistin exerts bactericidal activity by targeting LPS in the cytoplasmic membrane. We then exploited this information to devise a new therapeutic approach. Using the LPS transport inhibitor murepavadin, we were able to cause LPS accumulation in the cytoplasmic membrane, which resulted in increased susceptibility to colistin *in vitro* and improved treatment efficacy *in vivo*. These findings reveal new insight into the mechanism by which colistin kills bacteria, providing the foundations for novel approaches to enhance therapeutic outcomes.

## Introduction

The emergence of multi-drug-resistant Gram-negative pathogens such as *Escherichia coli, Klebsiella pneumoniae* and *Pseudomonas aeruginosa* has led to the increased use of polymyxin antibiotics, which are often the only viable last-resort therapeutic option [1–4]. Two closely related polymyxin antibiotics are used clinically, colistin (polymyxin E) and polymyxin B, which share a high degree of structural similarity, consisting of a cationic peptide ring of 7 amino acids connected to a hydrophobic acyl tail by a linear chain of 3 amino acids [1,3].

Polymyxins are rapidly bactericidal towards Gram-negative bacteria *in vitro* but are considerably less efficacious *in vivo*, with up to 70% of patients failing to respond to colistin treatment [5–7]. Restrictions on dosage due to the nephrotoxicity of polymyxins means that only 50% of people with normal renal function achieve a steady state serum concentration sufficient to kill bacteria [8,9]. As such, there is a desperate need to develop new approaches to enhance the efficacy of polymyxin antibiotics.

Barriers to increasing polymyxin efficacy include the significant gaps in our understanding of their mode of action. Whilst it is well established that the binding of polymyxins to LPS on the surface of Gram-negative bacteria leads to disruption of the outer membrane (OM), it is unclear how this results in cell lysis and bacterial death (Supplementary Fig. 1) [3,10]. It is hypothesised that damage to the LPS monolayer enables polymyxins to traverse the OM via a process of ‘self-directed uptake’, although this has not been demonstrated experimentally [10,11]. Once across the OM, polymyxins permeabilize the cytoplasmic membrane (CM), which is required for bacterial lysis and killing [1–3]. However, the mechanism by which colistin damages the CM is unclear [11,12]. It has been proposed that the surfactant activity of polymyxins, conferred by the positively charged peptide ring and hydrophobic tail, is sufficient to compromise the phospholipid bilayer of the CM via a detergent-like effect [1,3]. In support of this, polymyxins can interact with mammalian cell membranes, leading to changes in epithelial monolayer permeability [13]. Polymyxin antibiotics also have some inhibitory activity against the Gram-positive bacterium *Streptococcus pyogenes*, where the CM is formed of a phospholipid bilayer [14].

However, several lines of evidence call into doubt the ability of physiologically relevant concentrations of polymyxins to disrupt phospholipid bilayers. Firstly, the concentrations of polymyxin B required to disrupt mammalian epithelial cells or inhibit the growth of *S. pyogenes* (8-16 μg ml^-1^) are above typical serum concentrations of the antibiotic, and colistin at clinically-relevant concentrations displays no activity against other Gram-positive organisms such as *Staphylococcus aureus* and *Enterococcus faecalis* [13–16]. Furthermore, colistin has very little activity against synthetic phospholipid bilayer membranes unless LPS is present, a finding that explains why polymyxins are 30-100-fold less active against colistin-resistant *A. baumanii* isolates that are LPS-deficient, with an OM composed of a phospholipid bilayer [17–19]. Finally, molecular dynamics simulations show that the interaction of colistin with phospholipid bilayers is unlike what has been reported for other antimicrobial peptides that target phospholipid bilayers [20]. Together, these observations call into question whether, at physiologically relevant concentrations, colistin disrupts the CM of Gram-negative bacteria via the engagement of the polymyxin antibiotic with membrane phospholipids.

In addition to the mode of action of colistin, there are also gaps in our understanding of the mechanisms by which colistin resistance protects bacteria from polymyxin antibiotics. In Gram-negative bacteria, LPS is synthesised in the cytoplasm via the Raetz pathway, during which it is introduced into the inner leaflet of the CM [26,27]. It is then flipped to the outer leaflet of the CM by MsbA before transportation to the OM via the LptABCDEFG machinery [28–30]. To date, 10 mobile colistin resistance (*mcr*) gene variants have been described, all of which encode phosphoethanolamine (pEtN) transferases that modify the lipid A component of LPS with pEtN as it is trafficked through the CM on the way to the OM [21–25]. Colistin resistance can also arise via mutations in genes encoding two-component regulatory systems such as PhoPQ, PmrAB or BasRS [31,32]. This typically leads to the addition of 4-amino-4-deoxy-l-arabinose (L-ara4N) and/or pEtN groups to LPS, with this modification also occurring at the CM [27,31].

Despite the association between MCR-mediated LPS modification and colistin resistance, there is evidence that it does not prevent polymyxin-mediated damage of the OM. For example, colistin has been shown to permit ingress of the *N*-phenyl-1-napthylamine (NPN) fluorophore into the OM of *E. coli* expressing *mcr-1* [10]. Furthermore, colistin greatly enhances the activity of hydrophobic antibiotics such as rifampicin against polymyxin-resistant bacteria via disruption of the OM [33]. However, despite colistin damaging the OM of resistant bacteria, it is unable to kill or lyse them [10]. This suggests that the modification of LPS with pEtN and/or L-ara4N protects the CM from colistin, but it is not clear how [10,33].

Improving our knowledge of how colistin kills bacteria is essential to help devise new approaches to enhance the efficacy of last resort polymyxin antibiotics [21–25]. To do this, we set out to better understand how *mcr*-1 protects bacteria from colistin and to then use this information to elucidate the mode of action of colistin, with the ultimate aim of exploiting this information to improve colistin efficacy.

## Results

### MCR-1 protects the CM but not the OM from colistin-mediated disruption

The first issue we wanted to resolve was whether MCR-1 protected the CM and/or OM of bacteria from colistin. To do this, we used an isogenic *E. coli* MC1000 strain pair, one of which expresses *mcr-1* from the IPTG-inducible vector pDM1 (*mcr*-1) to ensure consistent expression under our experimental conditions, and the other transformed with the pDM1 vector alone as a control (pEmpty) [34]. As expected, we found that *E. coli* MC1000 expressing *mcr-1* had a significantly greater colistin minimum inhibitory concentration (MIC, 2 μg ml^-1^) compared to the MC1000 pEmpty control strain (0.25 μg ml^-1^), which was similar to that seen for clinical isolates [34] (Supplementary Fig. 2). This confirmed that the *E. coli* cells were producing functional MCR-1.

To fully characterise the LPS-modifying activity of MCR-1, we undertook MALDI-TOF-based lipidomic analysis of both whole *E. coli* cells and *E. coli* spheroplasts that lacked an OM [35]. We confirmed spheroplast formation by microscopy and used FITC labelling of OM surface proteins to demonstrate removal of the OM (Supplementary Fig. 3, 4). Our lipidomic analysis revealed the presence of LPS modified with pEtN in both the CM and OM of *mcr*-1 expressing bacteria, consistent with the location of MCR-1 in the CM [34,36] (Supplementary Fig. 5). Of note, whilst 42±19% of total cellular LPS from MCR-1-producing *E. coli* was unmodified, the proportion of unmodified LPS in the CM was just 21±2% (Fig. 1A, Supplementary Fig. 5).

**Figure 1.**
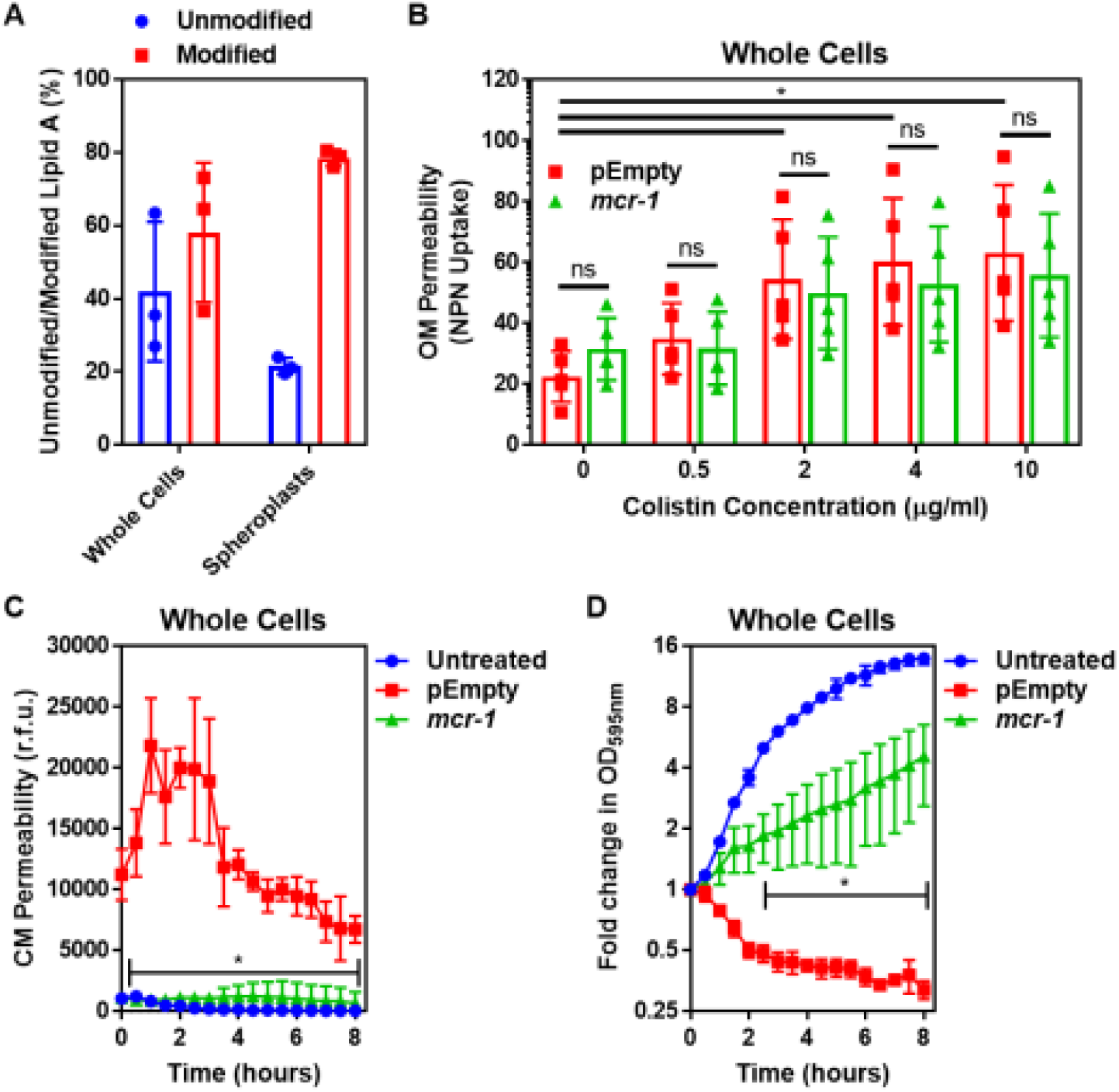
Colistin disrupts the outer membrane but not the cytoplasmic membrane of *E. coli* expressing *mcr-1*. **A,** Quantification of LPS modified with phosphoethanolamine, expressed as the percentage of unmodified lipid A and unmodified lipid A, in whole cells and spheroplasts of E. coli MC1000 expressing *mcr-1*, as determined by MALDI-TOF-based lipidomics (n=3 in duplicate, *p<0.05 between Whole Cells and Spheroplasts). **B,** OM disruption of *E. coli* MC1000 cells expressing *mcr-1* or an empty plasmid control strain (pEmpty) during 10 minutes of exposure to colistin at the indicated antibiotic concentrations, as determined by uptake of the fluorescent dye NPN (10 μM) (n=5, each data point represents the arithmetic mean of 20 replicate measurements; ns: p>0.05 between pEmpty and *mcr-1* strains, *p<0.05 between the indicated concentrations of colistin). **C,** Permeabilization of the CM of *E. coli* MC1000 cells expressing *mcr-1* or empty plasmid-containing cells during incubation with colistin (4 μg ml^-1^), as determined using 2.5 μM propidium iodide (PI) (n=4; *p<0.0001 between pEmpty and *mcr-1* strains). **D,** Growth or lysis of *E. coli* MC1000 cells expressing *mcr-1* or empty plasmid control cells during exposure to colistin (4 μg ml^-1^), as measured using OD_595nm_ readings (n=4; *p<0.05 between pEmpty and *mcr-1* strains). Data in **A** were analysed by a two-tailed paired Student’s t-test. Data in **B-D** were analysed by a two-way ANOVA with Sidak’s **(B)** or Dunnett’s **(C,D)** post-hoc tests. Data are presented as the arithmetic mean, and error bars represent the standard deviation of the mean. OM: outer membrane; NPN: *N*-phenyl-1-naphthylamine; CM: cytoplasmic membrane; r.f.u.: relative fluorescence units; OD: optical density.

We next assessed the effect of colistin on the integrity of the *E. coli* OM using the hydrophobic NPN dye, which fluoresces upon contact with phospholipids exposed by damage to the LPS monolayer [10,37]. As expected, colistin caused permeabilization of the OM of the *E. coli* pEmpty strain in a dose-dependent manner (Fig. 1B). In agreement with previous findings, we found that colistin also disrupted the OM of *E. coli* expressing *mcr*-1 to a similar degree to *E. coli* pEmpty (Fig. 1B) [10]. Therefore, MCR-1-mediated changes to LPS did not prevent permeabilization of the OM by colistin, which reflects the presence of the relatively large quantity of unmodified LPS in the OM as determined in our lipidomic analysis (Fig. 1A, Supplementary Fig. 5).

Next, we assessed damage to the CM structure in the *E. coli* strain pair during colistin exposure, using the membrane impermeant dye propidium iodide (PI). PI fluoresces upon contact with DNA in the bacterial cytoplasm, and thus is indicative of permeabilization of the both the OM and CM of whole bacterial cells [38,39]. As expected, colistin exposure resulted in a strong PI signal from *E. coli* pEmpty cells, indicative of CM permeabilization (Fig. 1C). However, despite colistin permeabilizing the OM of *E. coli* expressing *mcr*-1, the CM of these bacteria remained intact, as demonstrated by the lack of PI-mediated fluorescence (Fig. 1C). In keeping with these findings, colistin caused lysis of *E. coli* pEmpty cells, as seen by a reduction in OD_595nm_ readings over time (Fig. 1D). By contrast, *E. coli* cells producing MCR-1 grew in the presence of colistin despite the damage the polymyxin caused to the OM, as demonstrated by an increase in OD_595nm_ measurements over time (Fig. 1D). Taken together, these data demonstrate that MCR-1 protects the CM but not the OM from colistin.

### Colistin targets LPS in the CM

To explain the ability of MCR-1-mediated LPS modification to protect the CM from colistin, we considered two possibilities. Firstly, that LPS modifications alter the biophysical properties of the phospholipid bilayer of the CM [10]. Secondly, that colistin targets LPS in the CM and that modification with pEtN confers protection.

To test the first possibility, we determined the susceptibility of spheroplasts from *E. coli* MC1000 *mcr*-1 or pEmpty to colistin or the cationic antimicrobial peptides (CAMPs) daptomycin or nisin. Both CAMPs are well characterised for their ability to permeabilize phospholipid bilayers and, like colistin, they are positively charged, enabling us to detect whether any changes to membrane charge or fluidity conferred by MCR-1-modified LPS in the CM contributed to polymyxin resistance [40,41].

The CM of spheroplasts of *E. coli* MC1000 *mcr*-1 was resistant to damage by colistin, but susceptible to daptomycin and nisin (Fig. 2A,B,C). By contrast, colistin, daptomycin and nisin all permeabilized the CM of spheroplasts of *E. coli* pEmpty (Fig. 2A,B,C). In keeping with the data from assays measuring CM damage, colistin, daptomycin and nisin all caused lysis of spheroplasts of *E. coli* pEmpty, whilst the spheroplasts from *E. coli* expressing *mcr-1* were undamaged by colistin, but were lysed by both daptomycin and nisin (Fig. 2D,E,F). Combined, these data demonstrated that the protection afforded to the CM by MCR-1 is specific for colistin, and that that the polymyxin antibiotic does not share the same target as the phospholipid-targeting CAMPs. Thus, since MCR-1 specifically modifies LPS and selectively protects the CM from colistin, we concluded that LPS is the CM target of colistin, just as it is in the OM.

**Figure 2.**
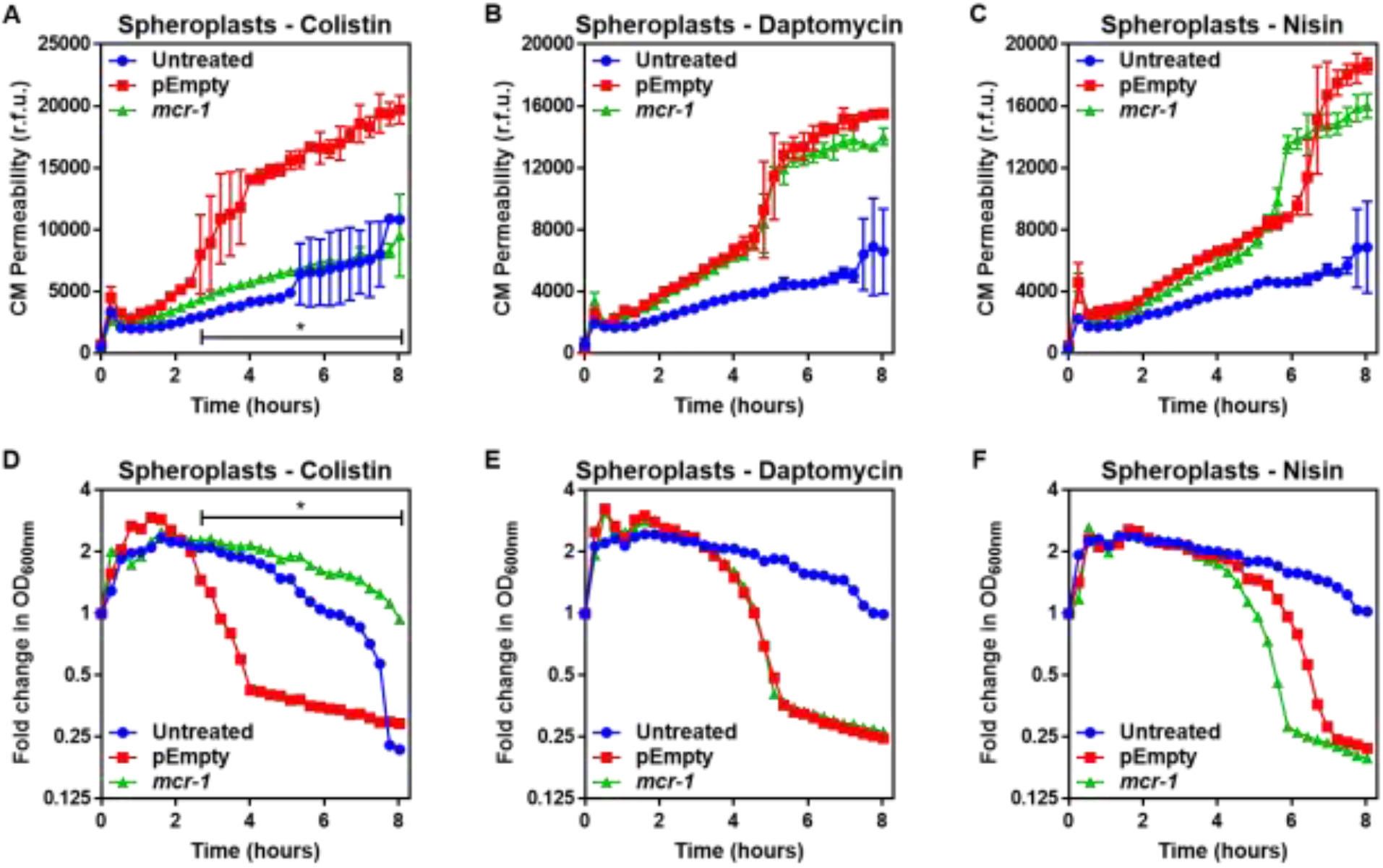
MCR-1 protects the cytoplasmic membrane of *E. coli* spheroplasts from colistin but not other cationic antimicrobial peptides. **A-C,** Permeabilization of the CM of *E. coli* MC1000 spheroplasts generated from bacteria expressing *mcr-1* or empty plasmid control bacteria (pEmpty) during incubation with **(A)** colistin (4 μg ml^-1^), **(B)** daptomycin (20 μg ml^-1^, with 1.25 μM Ca^2+^ ions) or **(C)** nisin (20 μg ml^-1^), as determined using 0.25 μM PI (n=3, experiment performed on 4 independent occasions; *p<0.01 between pEmpty and *mcr-1* strains). **D-F,** Lysis of *E. coli* MC1000 spheroplasts generated from bacteria expressing *mcr-1* or empty plasmid control bacteria during incubation with **(D)** colistin (4 μg ml^-1^), **(E)** daptomycin (20 μg ml^-1^, with 1.25 μM Ca^2+^ ions) or **(F)** nisin (20 μg ml^-1^), as measured using OD_600nm_ readings (n=3, experiment performed on 4 independent occasions; *p<0.05 between pEmpty and *mcr-1* strains, error bars are omitted for clarity). Data in **A-F** were analysed by a two-way ANOVA with Dunnett’s post-hoc test. Data are presented as the arithmetic mean, and error bars, where shown, represent the standard deviation of the mean. CM: cytoplasmic membrane; r.f.u.: relative fluorescence units; OD: optical density.

### Colistin damages the CM by disrupting cation bridges between LPS molecules

To understand how colistin targeting of LPS in the CM leads to membrane disruption, we studied the role of cation bridges which are crucial for stabilising interactions between LPS molecules, by exposing spheroplasts from *E. coli* pEmpty cells to colistin in the absence or presence of excess magnesium. In keeping with a role for cation bridges, we found that magnesium chloride conferred dose-dependent protection from colistin-mediated CM disruption (Fig. 3A). To rule out a general protective osmotic effect from the higher salt concentration, we demonstrated that identical concentrations of sodium chloride did not protect spheroplasts from colistin (Fig. 3B). Furthermore, the presence of exogenous magnesium had no significant effect on reducing spheroplast CM damage caused by daptomycin or nisin (Fig. 3CD), confirming that these CAMPs do not have the same CM target as colistin.

**Figure 3.**
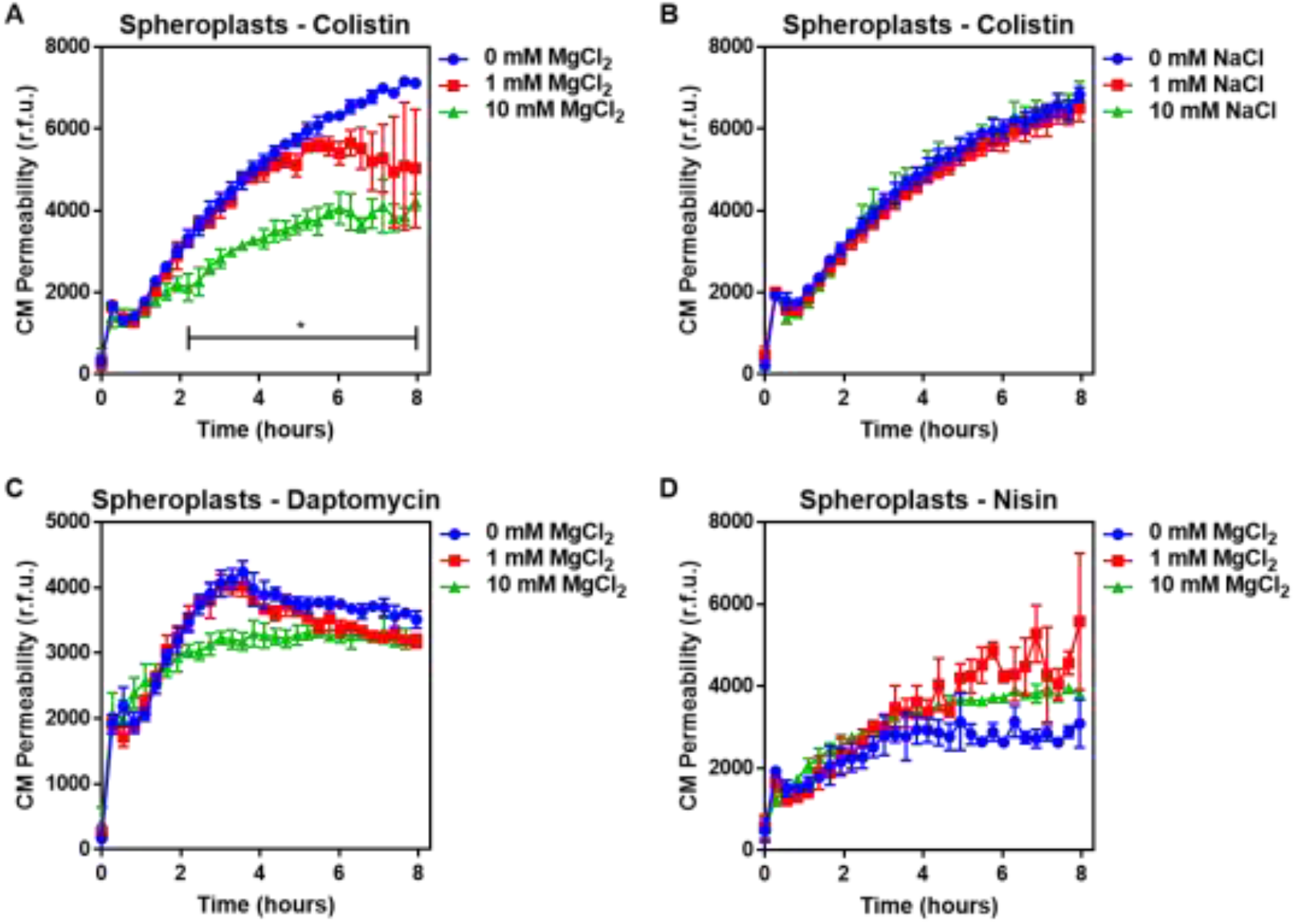
Colistin damages the cytoplasmic membrane by disrupting cation bridges between LPS molecules. **A, B,** Permeabilization of the CM of *E. coli* MC1000 spheroplasts generated from empty plasmid control bacteria during incubation with colistin (4 μg ml^-1^), in the absence or presence of either MgCl2 **(A)** or NaCl **(B)** at the indicated concentrations, as determined using 0.25 μM PI (n=3, experiment performed on 3 independent occasions; *p<0.01 between 0 mM MgCl2 and 10 mM MgCl2). **C, D,** Permeabilization of the CM of *E. coli* MC1000 spheroplasts generated from empty plasmid control bacteria during incubation with either **(C)** daptomycin (20 μg ml^-1^, with 1.25 μM Ca^2+^ ions) or **(D)** nisin (20 μg ml^-1^), in the absence or presence of MgCl2 at the indicated concentrations, as determined using 0.25 μM PI (n=3, experiment performed on 3 independent occasions). Data in **A-D** were analysed by a two-way ANOVA with Dunnett’s post-hoc test. Data are presented as the arithmetic mean, and error bars represent the standard deviation of the mean. CM: cytoplasmic membrane; r.f.u.: relative fluorescence units.

In conclusion, our findings demonstrate that colistin targets LPS in the CM of polymyxin-susceptible *E. coli*, leading to the displacement of cationic inter-LPS bridges, membrane disruption and ultimately bacterial lysis. This is similar to the mechanism by which colistin disrupts the OM of bacteria and synthetic phospholipid bilayer membranes containing low levels of LPS [17,44–45]. However, the high levels of LPS modified by pEtN in the CM of MCR-1-producing *E. coli* prevent colistin from targeting LPS in the CM, protecting the membrane and conferring resistance to the polymyxin antibiotic.

### Murepavadin-triggered LPS accumulation in the cytoplasmic membrane sensitises *P. aeruginosa* to colistin

Having determined that colistin kills bacteria by targeting LPS in the CM, we wanted to use this information to develop a new therapeutic approach to enhance colistin efficacy.

Murepavadin is a first in class peptide-based inhibitor of the LptD component of the LptABCDEFG complex of *P. aeruginosa* that transports LPS from the CM to the OM [46]. Thus, inhibition of the Lpt system in *P. aeruginosa* leads to LPS accumulation in the CM, which we hypothesised would increase the susceptibility of the bacterium to colistin [46,47].

To test our hypothesis, we first used a checkerboard MIC assay and found that colistin synergised with murepavadin against *P. aeruginosa* PA14 cells, revealing that sub-lethal concentrations of the LptD inhibitor sensitised the bacterium to colistin (Fig. 4A) [48]. To confirm that sub-lethal concentrations of murepavadin altered LPS abundance in the CM, *P. aeruginosa* was incubated with murepavadin, before the amount of LPS in whole cells and spheroplasts was measured using the well-established Limulus Amoebocyte Lysate (LAL) assay (Supplementary Fig. 6, 7) since previous work has shown this approach to be a highly accurate way of quantifying LPS in whole cell lysates [49]. The suitability of the LAL assay was further confirmed using a MALDI-TOF-based lipidomic analysis of spheroplast lysates, which confirmed that the LPS in both murepavadin-exposed and untreated bacteria was unmodified and thus able to be accurately detected and quantified (Supplementary Fig. 8) [50]. Sub-lethal concentrations of murepavadin caused a slight reduction in LPS levels in the OM of *P. aeruginosa* cells, but a significant increase in the amount of LPS in the CM compared to untreated spheroplasts (Supplementary Fig. 8, 9). Moreover, our lipidomic analysis revealed that the ratio of lipid A:phospholipid increased in *P. aeruginosa* spheroplasts pre-exposed to murepavadin, confirming that the LptD inhibitor caused LPS to accumulate in the CM (Supplementary Fig. 8).

**Figure 4.**
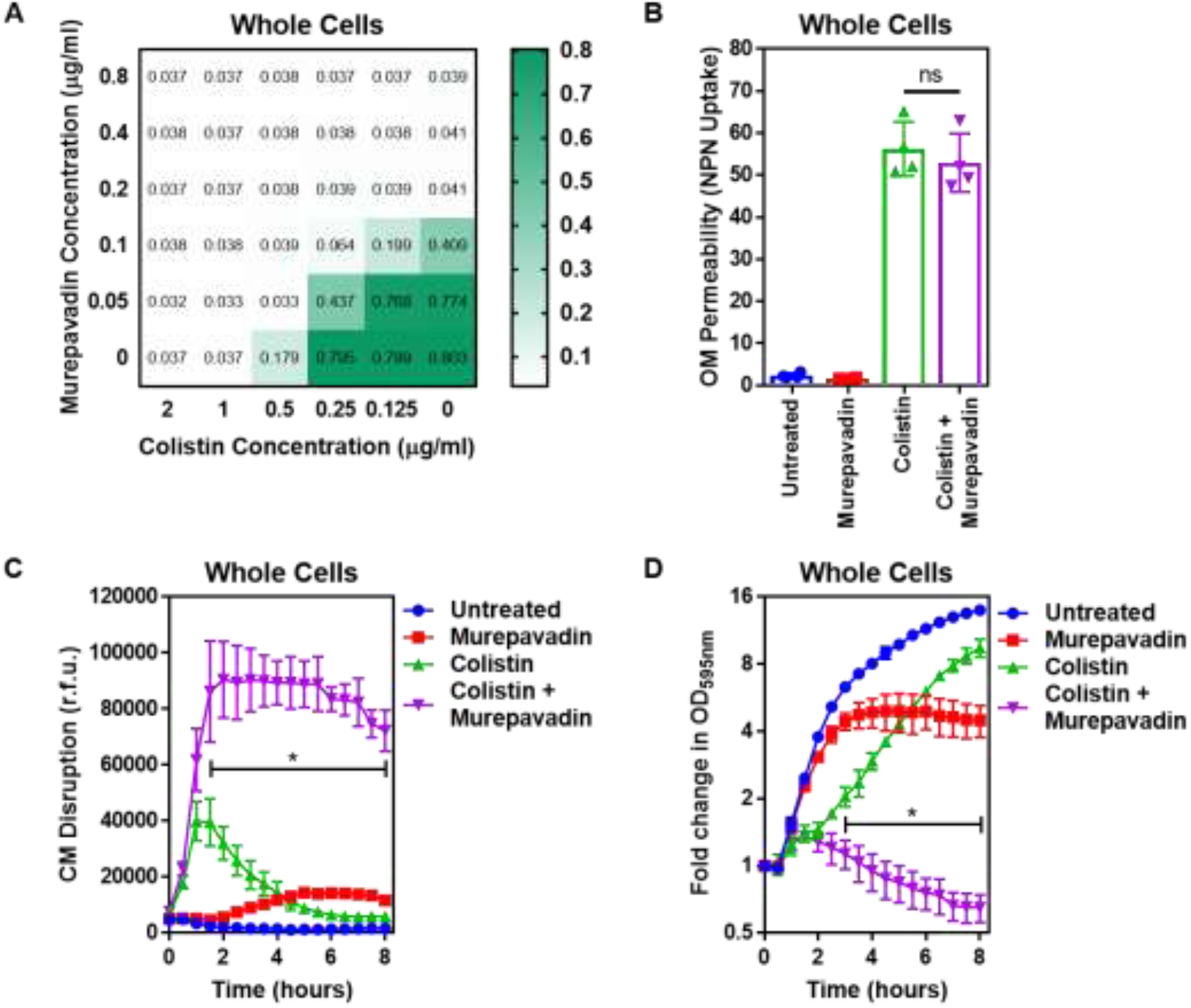
Murepavadin sensitises *P. aeruginosa* to colistin by increasing LPS abundance in the cytoplasmic membrane. **A,** Checkerboard broth microdilution assay showing the synergistic growth-inhibitory interaction between colistin and the LPS transport inhibitor murepavadin against *P. aeruginosa* PA14 cells, as determined by measuring OD_595nm_ after 18 hours incubation. **B,** OM disruption of *P. aeruginosa* PA14 cells during 10 minutes exposure to colistin (2 μg ml^-1^) in the absence or presence of murepavadin (0.05 μg ml^-1^), as assessed by uptake of the fluorescent dye NPN (10 μM) (n=4, each data point represents the arithmetic mean of 20 replicate measurements; ns: p>0.05 between colistin-treated bacteria). **C,** CM disruption of *P. aeruginosa* PA14 cells exposed to colistin (2 μg ml^-1^) in the absence of presence of murepavadin (0.05 μg ml^-1^), as determined using 2.5 μM PI (n=4; *p<0.0001 for Colistin and Murepavadin-exposed cells compared to Colistin alone). **D,** Lysis of *P. aeruginosa* PA14 cells exposed to colistin (2 μg ml^-1^) in the absence of presence of murepavadin (0.05 μg ml^-1^), as measured by OD_595nm_ readings (n=4; *p<0.01 for Colistin and Murepavadin-exposed cells compared to Colistin alone). Data in **B** were analysed by a one-way ANOVA with Tukey’s post-hoc test. Data in **C-D** were analysed by a two-way ANOVA with Dunnett’s post-hoc test. Data are presented as the arithmetic mean, and error bars represent the standard deviation of the mean. OM: outer membrane; NPN: *N*-phenyl-1-naphthylamine; CM: cytoplasmic membrane; r.f.u.: relative fluorescence units; OD: optical density.

Next, we proceeded to test whether LPS accumulation in the CM increased the susceptibility of *P. aeruginosa* to colistin. We started by examining the effect of colistin on the OM and CM of *P. aeruginosa* exposed, or not, to murepavadin. Despite the slight reduction of LPS at the OM caused by murepavadin, colistin permeabilized the OM to the same extent as bacteria that had not been exposed to murepavadin, with similar levels of NPN uptake (Fig. 4B). By contrast, however, murepavadin significantly enhanced permeabilization of the CM by colistin in whole cells of *P. aeruginosa*, as determined via uptake of PI (Fig. 4C). Thus, an increase in LPS levels in the CM promoted colistin-mediated damage, in keeping with our conclusion that LPS in the CM is the target of the polymyxin antibiotic. Furthermore, *P. aeruginosa* cells exposed to murepavadin were more rapidly lysed by colistin than untreated cells, confirming that LPS accumulation in the CM increased susceptibility to the polymyxin antibiotic (Fig. 4D).

### Combination therapy with colistin and murepavadin promotes clearance of *P. aeruginosa*

Having shown that murepavadin sensitised the CM to colistin-mediated damage, we wanted to determine whether this translated into enhanced antibacterial activity against relevant clinical isolates and increased treatment efficacy *in vivo*. We found that a sub-lethal concentration of murepavadin sensitised *P. aeruginosa* PA14 to a normally sub-lethal concentration of colistin (2 μg ml^-1^), resulting in >10,000-fold reduction in c.f.u. counts relative to bacteria incubated with murepavadin or colistin alone after 8 h (Fig. 5A). We also found that murepavadin potentiated the activity of an even lower concentrations of colistin (1 μg ml^-1^), with exposure to the LPS transport inhibitor increasing the ability of the polymyxin antibiotic to damage the CM, triggering bacterial lysis and cell death (Supplementary Fig. 10).

**Figure 5.**
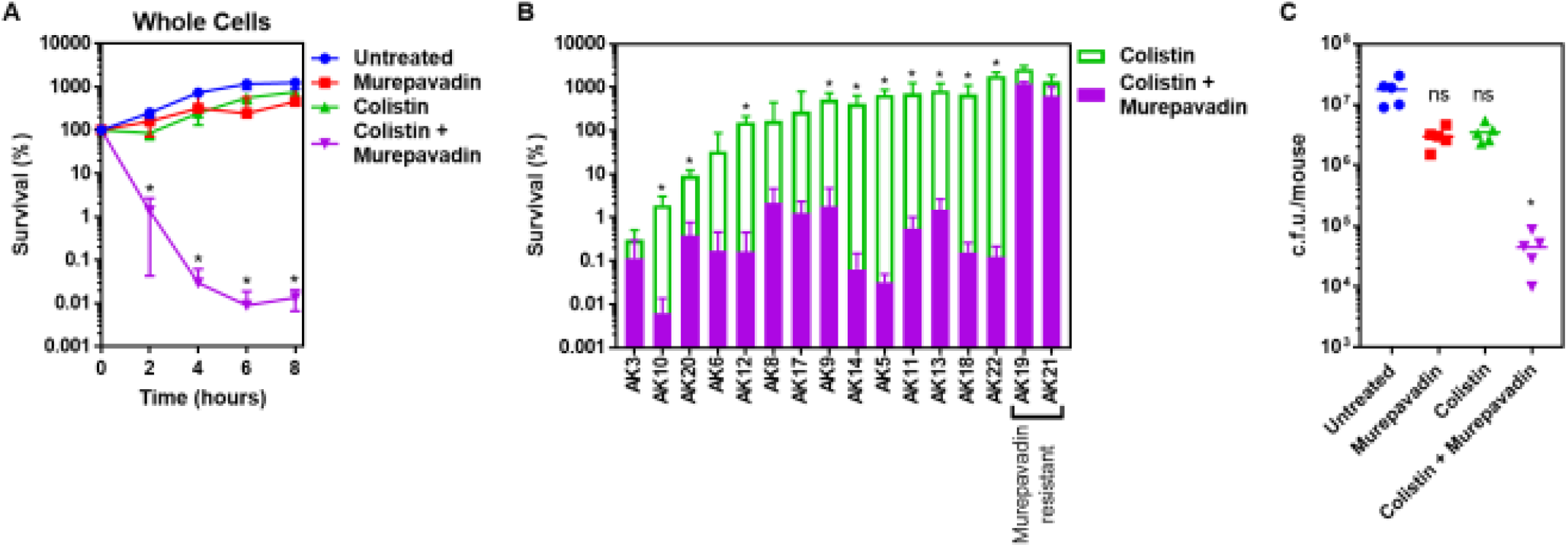
Colistin-murepavadin combination therapy promotes killing of *P. aeruginosa in vitro* and *in vivo*. **A,** Survival of *P. aeruginosa* PA14 cells exposed to colistin (2 μg ml^-1^) in the absence of presence of murepavadin (0.05 μg ml^-1^), as determined by c.f.u. counts (n=4; *p<0.05 for Colistin and Murepavadin-exposed cells compared to Colistin alone). **B,** Survival or growth of a panel of clinical multidrug-resistant *P. aeruginosa* strains isolated from the sputum of cystic fibrosis patients after 8 hours exposure to colistin (2 μg ml^-1^) alone, or in the presence of a sub-lethal concentration (0.5x MIC) of murepavadin, as determined by c.f.u. counts (n=4; *p<0.01 for Colistin and Murepavadin-exposed cells compared to Colistin alone). **C,** Burden of *P. aeruginosa* PA14 in the lungs of mice after 3 hours treatment with murepavadin (0.25 mg/kg), colistin (5 mg/kg), neither antibiotic, or both antibiotics in combination, as determined by c.f.u. counts (each data point represents a single mouse; for each group, n=5; ns: p>0.05, *p<0.001 compared to Untreated mice). Data in **A-B** were analysed by a two-way ANOVA with Dunnett’s **(A)** or Sidak’s **(B)** post-hoc tests. Data in **C** were analysed by a Kruskal-Wallis test with Dunn’s post-hoc test. Data are presented as the arithmetic mean, and error bars, where shown, represent the standard deviation of the mean.

We next examined a panel of 16 multi-drug resistant *P. aeruginosa* clinical strains, isolated from the sputum of cystic fibrosis (CF) patients, to investigate whether murepavadin increased colistin-mediated bacterial killing (Supplementary Fig. 11). Of these 16 clinical isolates, 14 were susceptible to murepavadin alone, whilst two strains were resistant to the LptD inhibitor (Supplementary Table 2). In 11 out of 14 murepavadin-susceptible CF isolates tested (79%), sub-lethal concentrations of murepavadin caused a significant increase in the bactericidal activity of colistin against *P. aeruginosa* (Fig. 5B). Importantly, murepavadin did not affect the bactericidal activity of colistin against the two strains that were resistant to LptD inhibitor (Fig. 5B). This confirmed that the potentiating effects of the LptD inhibitor on polymyxin-mediated killing were not due to off-target effects.

Next, we wanted to test whether colistin-murepavadin dual therapy could promote bacterial clearance during infection *in vivo*. We inoculated mice via the intranasal route with *P. aeruginosa* PA14 to cause a lung infection, and then treated intranasally with PBS alone, or PBS containing colistin only (5 mg kg^-1^), murepavadin only (0.25 mg kg^-1^), or colistin and murepavadin combined at the concentrations used for mono-treatment. These concentrations were based on those used previously to mimic treatment of human lung infections, and the route of delivery is similar to that used clinically [51–54]. Mono-therapy with colistin alone or murepavadin alone had very little effect on the bacterial load assessed after 3 hours treatment compared with the no-treatment control (Fig. 5C). By contrast, combination therapy with colistin and murepavadin caused a ~500-fold reduction in c.f.u. counts relative to the no-treatment control (Fig. 5C). Therefore, murepavadin synergises with colistin both *in vitro* and *in vivo*, suggesting it may be useful as a combination therapeutic approach for lung infections caused by *P. aeruginosa*.

## Discussion

Colistin is an increasingly important last-resort antibiotic used to treat infections caused by multi-drug-resistant Gram-negative pathogens, including *P. aeruginosa, K. pneumoniae* and *E. coli* [2–4]. However, treatment failure occurs frequently, and resistance is a growing concern [5–10]. Efforts to address these issues are compromised by a poor understanding of colistin’s bactericidal mode of action. Whilst the initial interactions of colistin with LPS in the OM of Gram-negative bacteria were well-established, it was unclear how the antibiotic traversed the OM and damaged the CM to cause cell lysis (Supplementary Fig. 1). In this work, we demonstrate that colistin targets LPS in the CM, resulting in membrane permeabilization, bacterial lysis and killing (Fig. 6).

**Figure 6.**
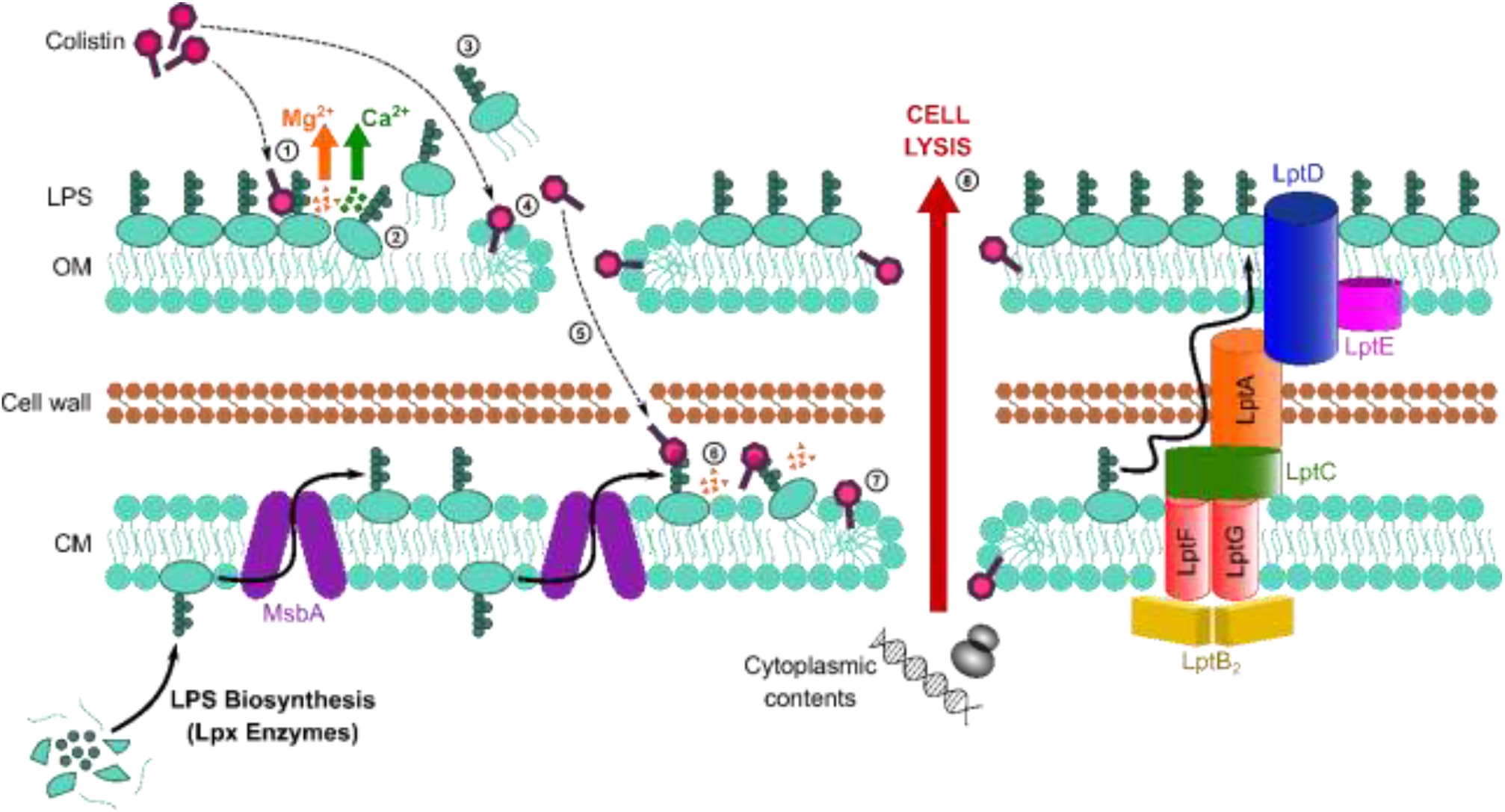
Colistin kills bacteria by targeting LPS in both the outer and cytoplasmic membranes, leading to disruption of the cell envelope and bacterial lysis. Diagrammatic representation of the novel proposed mechanism of action of colistin: Colistin binds to LPS in the OM (1), displacing cations that form bridges between LPS molecules, which leads to destabilisation of the OM (2). As a consequence of the weakening of intermolecular bonds in the LPS monolayer, LPS is released from the bacterial surface (3), allowing colistin to further damage the OM via the action of the polymyxin lipid tail (4). This provides colistin with access to the periplasm, where colistin interacts with LPS in the CM (5) that is awaiting transport to the OM by the LptABCDEFG machinery after being synthesized in the cytoplasm and flipped to the outer leaflet of the CM by MsbA. As in the OM, colistin binding to LPS results in displacement of cation bridges and disruption of the CM (6), which it ultimately permeabilizes (7), culminating in the loss of cytoplasmic contents, cell lysis and bacterial death (8).

Our conclusion that colistin targets LPS in the CM was based initially on experiments with *E. coli* expressing the *mcr*-1 colistin resistance determinant. MCR-1 modifies lipid A with a phosphoethanolamine moiety as it passes through the CM on its way to the OM [21,23]. Since MCR-1 specifically protected spheroplasts from colistin but not nisin or daptomycin, which both target phospholipid bilayers, it was clear that colistin did not share the same target as the other two CAMPs. Given the only difference between *E. coli* spheroplasts expressing *mcr*-1 and pEmpty spheroplasts was modified LPS, our data reveal that colistin targets LPS in the CM, leading to disruption of the CM, which is a pre-requisite for subsequent cell lysis and bacterial killing [38]. These findings were then supported by experiments showing that LPS accumulation in the CM of *P. aeruginosa* sensitised this bacterium to colistin.

Whilst the abundance of LPS in the CM is 10-20 times lower than the OM, studies with model membranes have shown that the presence of LPS at a 1:100 ratio with phospholipids was both necessary and sufficient for colistin-mediated permeabilization [17,42–43]. Therefore, our conclusion explains how an antibiotic with a high degree of specificity for LPS could damage both the OM and CM [1]. The reason why MCR-1 protected the CM but not OM from colistin-mediated damage is most likely due to the higher proportion of unmodified LPS at the CM (21±2%) relative to the OM (42±19%) (Fig. 1A). Furthermore, the overall abundance of LPS in the CM is very low, resulting in very few targets (i.e. unmodified LPS molecules) for colistin in the CM of *mcr-1*-expressing *E. coli*. By contrast, the OM of MCR-1-producing *E. coli* contains many more unmodified LPS molecules that can be bound by colistin, explaining why colistin is able to damage this structure, but cannot permeabilize the CM of *E. coli* expressing *mcr-1* at a physiologically relevant concentration [42,43]. Whether colistin resistance conferred by chromosomal mutations in two-component systems is also mediated by modified LPS in the CM remains to be tested [22–27].

Our data showing that colistin requires unmodified LPS to be present in the CM in order to kill bacteria explains how an antibiotic with high affinity and specificity for LPS causes disruption to both the OM and CM [1]. Furthermore, these findings provide support for the observations that colistin does not damage the OM of colistin-resistant *A. baumannii* isolates where LPS has been replaced by a phospholipid bilayer, and that polymyxins cause only minimal disruption to model phospholipid membranes unless LPS is present [17–20].

Whilst the interaction of colistin with LPS in the CM is likely to share similarities with the same process at the OM, there are also likely to be differences owing to the differing concentrations of LPS between the two membranes [17,42–43]. In the OM, LPS is a highly abundant component with molecules tightly packed and stabilised with cation bridges. By contrast, LPS is a minority component in the CM, which may affect the rate and degree to which the CM is disrupted by polymyxins. In support of this, whilst colistin induced OM damage within minutes of bacterial exposure to the antibiotic, disruption of the CM took much longer. Even when spheroplasts lacking an OM were exposed to colistin, it still took more than 2 hours for CM permeabilization to occur. Therefore, it appears that colistin-mediated disruption of the CM is considerably less efficient than that of the OM, likely due to the much lower levels of LPS present in the CM.

Combined, these findings provide strong evidence that colistin targets LPS in the CM, in addition to the OM, and that this is required for the bactericidal and lytic activity of the antibiotic at clinically relevant concentrations. This insight into the mode of action of colistin enabled us to devise a new therapeutic approach to enhance colistin efficacy. Using the LptD inhibitor murepavadin, which is in development as an inhaled treatment for *P. aeruginosa* infections, we triggered LPS accumulation in the CM, and thereby increased the susceptibility of bacteria to colistin. The potential clinical utility of this approach was demonstrated by showing enhanced activity of colistin-murepavadin combination therapy against a panel of clinical CF isolates, as well as potent efficacy in a murine model of *P. aeruginosa* lung infection. It is anticipated that a combination of colistin and murepavadin could enhance the low treatment efficacy of polymyxin antibiotics and may also limit the toxic side effects associated with both compounds by enabling the use of lower doses of the drugs [55].

In summary, this work contributes to our understanding of the mechanism of action of colistin by demonstrating that polymyxin antibiotics target LPS in both the OM and the CM, and that this leads to the disruption of both membranes, resulting in the bactericidal and lytic activities of the antibiotic. Modulation of LPS levels can enhance colistin activity, providing the foundations for new approaches to enhance the efficacy of this antibiotic of last resort.

## Methods

### Bacterial strains and growth conditions

The bacterial strains used in this study are listed in Supplementary Table 1. For each experiment, all strains were grown in Luria broth (LB; Thermo Fisher Scientific, USA) for 18 h to stationary phase at 37°C with shaking (180 r.p.m.). For routine culture of bacteria on solid media, strains were grown on LB supplemented with 1.5% technical agar (BD Biosciences, USA). Liquid and solid growth media were supplemented with tetracycline (12.5 μg ml^-1^; Sigma-Aldrich, USA) and 0.5 mM isopropyl-β-D-thiogalactoside (IPTG, Melford Laboratories, UK) where required. Enumeration of bacterial c.f.u. was done by plating 10-fold serial dilutions of bacterial cultures on to Mueller-Hinton agar (MHA; Thermo Fisher Scientific) plates. Inoculated agar plates were incubated statically for 18 h in air at 37°C.

### Determination of minimum inhibitory concentrations of antibiotics

The minimum inhibitory concentration (MIC) of colistin and murepavadin against bacterial strains was determined by the broth microdilution protocol [56]. A microtitre plate was used to prepare a range of antibiotic concentrations in 200 μl cation-adjusted Mueller-Hinton broth (CA-MHB; Thermo Fisher Scientific) by two-fold serial dilutions. For certain experiments, checkerboard analyses were performed by preparing two-fold serial dilutions of two antibiotics in different directions, generating an 8 x 8 matrix to assess the MICs of the antibiotics in combination [48]. Stationary-phase bacteria were diluted 1000-fold in fresh CA-MHB and seeded into each well of the microtitre plate to a final concentration of 5 x 10^5^ c.f.u. ml^-1^. The microtitre plates were then incubated statically at 37°C for 18 h in air, after which point the MIC was defined as the lowest antibiotic concentration at which there was no visible growth of bacteria. In some cases, the extent of bacterial growth after 18 h incubation was also determined by obtaining OD_595nm_ measurements using a Bio-Rad iMark microplate absorbance reader (Bio-Rad Laboratories, USA).

### Bacterial growth assay

Stationary-phase bacteria were diluted 1000-fold in fresh CA-MHB, and 4 μl was seeded into the wells of a microtitre plate containing 200 μl CA-MHB, and for some experiments the LptD inhibitor murepavadin, to give a final inoculum of 5 x 10^5^ c.f.u. ml^-1^. The microtitre plate was incubated with shaking (180 r.p.m.) at 37°C for 16 h in a Tecan Infinite 200 Pro multiwell plate reader (Tecan Group Ltd., Switzerland) and optical density measurements were taken at 600 nm every 15 min.

### Production of spheroplasts

Spheroplasts of *E. coli* and *P. aeruginosa* strains lacking an OM and cell wall were generated as previously described [35]. Briefly, stationary-phase bacteria grown overnight were washed twice by centrifugation (12,300 x *g*, 3 min) followed by resuspension in CA-MHB, and added at a final inoculum of 10^8^ c.f.u. ml^-1^ to 9 ml CA-MHB containing for some experiments the LPS transport inhibitor murepavadin. Cultures were then incubated at 37°C with shaking (180 r.p.m.) for 2 h. After the incubation, bacteria were washed twice by centrifuging (3,273 x *g*, 20 min, 4°C) and resuspending first in 10 ml Tris buffer (0.03 M, pH 8.0; Sigma-Aldrich), and subsequently in Tris buffer (0.03 M, pH 8.0) containing 20% sucrose. EDTA (250 μl, 10 mg ml^-1^; Sigma-Aldrich) and lysozyme (1 ml, 10 mg ml^-1^; Roche, Switzerland) were added to remove the OM and cell wall respectively, and the cell suspension was incubated for 1 h in a water bath shaker at 30°C. Trypsin (500 μl, 10 mg ml^-1^; Sigma-Aldrich) was then added, and the culture again incubated at 30°C in a water bath shaker for 15 min. The resulting spheroplasts produced were harvested by mild centrifugation (2,000 x *g*, 20 min, 4°C), with the supernatant containing the removed OM extracted for further analysis. Successful conversion of bacterial whole cells into spheroplasts was confirmed using phase-contrast microscopy, as detailed below.

### Confirmation of successful spheroplast formation

Whole cells of *E. coli* and *P. aeruginosa* grown overnight were washed twice by centrifugation (12,300 x *g*, 3 min) and resuspension in CA-MHB, added at a final inoculum of 10^8^ c.f.u. ml^-1^ to 9 ml CA-MHB, and incubated for 2 h at 37°C with shaking (180 r.p.m.). Outer membrane proteins of these bacteria were subsequently labelled with fluorescein isothiocyanate (FITC, Sigma-Aldrich) as previously described [57]. Bacterial cells were washed twice by centrifugation (3,273 x *g*, 20 min, 4°C) and resuspension in 10 ml Labelling Buffer (50mM Na2CO3, 100 mM NaCl, pH 8.0), to which FITC was added at a final concentration of 0.5 mg ml^-1^. Bacteria were incubated for 30 mins at room temperature, before labelled cells were harvested by centrifuging (3,273 x *g*, 20 min, 4°C) and washed thrice by resuspending in 10 ml Tris buffer (0.03 M, pH 8.0) containing 20% sucrose. 1 ml of FITC-labelled bacteria was extracted and centrifuged (12,300 x *g*, 3 min), and the cells were fixed in 4% paraformaldehyde (PFA) in phosphate-buffered saline (PBS). The remaining 9 ml of FITC-labelled cells were converted into spheroplasts, as described above. The spheroplasts produced were recovered by mild centrifugation (2,000 x *g*, 20 min, 4°C) and resuspension in 9 ml Tris buffer (0.03 M, pH 8.0) containing 20% sucrose, before 1 ml of spheroplasts were fixed in the same way as with whole cells. The amount of FITC fluorescence in the OM of whole cells and CM of spheroplasts was observed using fluorescence microscopy, as described below. For quantification of FITC fluorescence, 200 μl samples of the fixed bacterial suspensions were seeded into the wells of a black-walled microtitre plate, and fluorescence measured with a Tecan Infinite 200 Pro multiwell plate reader, using an excitation wavelength of 490 nm and an emission wavelength of 525 nm.

### Microscopy

For phase-contrast and fluorescence microscopy, a 5 μl sample of fixed bacterial whole cells or spheroplasts was spotted onto a thin 1.2% agarose gel patch prepared in distilled water on a microscope slide. Bacteria were imaged using an Axio Imager.A2 Zeiss microscope (Carl Zeiss Microscopy GmbH, Germany) at 1000x magnification with an oil immersion objective lens. The ZEN 2012 software was used for image acquisition, whilst analysis of cell length:width ratios was done using the FIJI/ImageJ software by measuring two perpendicular lines drawn through the centre of bacteria. For each experiment, all microscopy images were acquired and processed using identical settings throughout.

### Determination of LPS concentration and modification by mass spectroscopy

Spheroplasts from bacterial cells were prepared as described above and then resuspended in ddH2O (200 μl), before mild acid hydrolysis was performed via the addition of 2% (vol/vol) acetic acid in ddH2O (200 μl) and incubation at 100°C for 30 min. For experiments with whole cells, bacteria grown overnight to stationary-phase were washed three times by centrifuging and resuspending in ddH2O, and a mild acid hydrolysis was performed on these whole cells as described for spheroplasts. Acid-treated whole cells or spheroplasts were recovered by centrifugation (17,000 × *g*, 2 min), and the resulting pellet was washed before being resuspended in 50 μl ultrapure water. The whole cell or spheroplast suspension (0.5 μl) was then loaded immediately onto the target and overlaid with 1.2 μl of a matrix consisting of 9H-Pyrido[3,4-B]indole (Norharmane) (Sigma-Aldrich) dissolved in 90:10 (vol/vol) chloroform/methanol to a final concentration of 10 mg ml^-1^. The bacterial suspension and matrix were then mixed on the target before gentle drying under air at room temperature. MALDI-TOF mass spectroscopy analysis was undertaken with a MALDI Biotyper Sirius system (Bruker Daltonics, USA), using the linear negative-ion mode as described previously [36]. Manual peak picking at masses relevant to phospholipids or lipid A was performed on the mass spectra obtained, and the corresponding signal intensities at the defined masses were determined. Peaks were considered only if their signal/noise ratio was at least 5. To determine the ratio of modified lipid A to unmodified lipid A, the area under the pETN-modified lipid A peak (*m/z* 1,919.2) was divided by the area under the peak corresponding to native lipid A (*m/z* 1,796.2). To determine the relative abundance of LPS, the sum of the area under the lipid A peaks (*m/z* 1447-1700) was divided by the sum of the area under representative phospholipid peaks (phospholipid 34:1,2, *m/z* 717-747). All mass spectra were generated and analysed with three biological replicates and two technical replicates.

### Outer membrane disruption assay

To detect damage to the OM of bacteria, the well-established NPN uptake assay was used [37]. Stationary-phase bacterial cells were washed in fresh CA-MHB and diluted to an optical density (OD_600nm_) of 0.5 in 5 mM pH 7.2 HEPES buffer (Sigma-Aldrich). This bacterial suspension was added to wells containing the relevant antibiotics in HEPES buffer, as well as the fluorescent probe *N*-phenyl-1-naphthylamine (NPN; Acros Organics, USA) at a final concentration of 10 μM. Samples were placed in a black microtitre plate with clear-bottomed wells and fluorescence measured immediately in a Tecan Infinite 200 Pro multiwell plate using an excitation wavelength of 355 nm and an emission wavelength of 405 nm. Fluorescence measurements were obtained every 30 seconds for 10 min, and the degree of OM permeabilization, referred to as the NPN Uptake Factor, was calculated using the following formula:

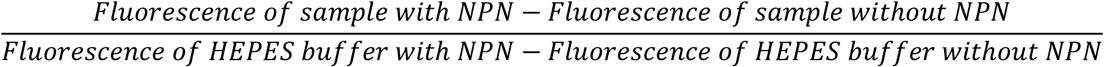

### Cytoplasmic membrane disruption assay

To measure CM disruption of whole cells, bacteria grown to stationary-phase overnight were washed and inoculated into 3 ml MHB containing the relevant antibiotics. Cultures were incubated at 37°C with shaking (180 r.p.m.) for up to 8 h, and every 30 min, aliquots (200 μl) were taken and bacteria isolated by centrifugation (12,300 x *g*, 3 min). Cells were then washed in sterile PBS before being added to the wells of a black-walled microtitre plate, and propidium iodide (PI, Sigma-Aldrich) was added to each well at a final concentration of 2.5 μM. Fluorescence was measured immediately in a Tecan Infinite 200 Pro multiwell plate reader (excitation at 535 nm, emission at 617 nm). To measure disruption of the CM in spheroplasts, spheroplasts of *E. coli* generated as detailed above were washed by centrifugation (4,000 x *g*, 5 min) and resuspension in Tris buffer (0.03 M, pH 8.0) containing 20% sucrose. Spheroplast samples (20 μl) were then added in the wells of a black-walled microtitre plate to 180 μl of Tris buffer (0.03 M, pH 8.0) containing 20% sucrose, the relevant antibiotics, and PI at a final concentration of 0.25 μM. The microtitre plate was incubated with shaking (180 r.p.m.) at 37°C for up to 8 h in a Tecan Infinite 200 Pro multiwell plate reader and fluorescence (excitation at 535 nm, emission at 617 nm) measured every 15 min. For both whole bacterial cells and spheroplasts, to account for differences in fluorescence values arising from variations in cell number, relative fluorescence unit (r.f.u.) measurements were corrected for OD at 600 nm.

### Determination of bacterial lysis

In the case of whole bacterial cells, washed stationary-phase bacteria were inoculated into 3 ml CA-MHB containing the relevant antibiotics, as described above. Cultures were then placed in a shaking incubator (37°C, 180 r.p.m.) for 8 h, and every 30 min, samples (200 μl) were transferred to a microtitre plate, where OD_595nm_ measurements were obtained using a Bio-Rad iMark microplate absorbance reader. For spheroplasts, washed spheroplasts (20 μl) were added to 180 μl of Tris buffer (0.03 M, pH 8.0) containing 20% sucrose and the relevant antibiotics in a microtitre plate as detailed above. The microtitre plate was incubated for up to 8 h at 37°C with shaking (180 r.p.m.) in a Tecan Infinite 200 Pro multiwell plate reader, and readings of OD_600nm_ were made every 15 min.

### Determination of LPS concentration by Limulus Amebocyte Lysate assay

Stationary-phase bacteria grown overnight were washed and grown for 2 h, before conversion to spheroplasts as described above. During formation of spheroplasts, the OM extracted from the bacterial cells was recovered, and the concentration of LPS in the OM, as well as the concentration of LPS in the CM of spheroplasts, was determined using the chromogenic Limulus Amebocyte Lysate (LAL) assay (all reagents from Thermo Fisher Scientific) as described previously [49–50, 58]. OM samples and spheroplasts lysed by freeze-thaw to release LPS were diluted in 10-fold steps, and 50 μl of each sample was equilibrated to 37°C and loaded into the wells of a microtitre plate at the same temperature. LAL reagent (50 μl) was added to each well, and the mixture incubated at 37°C for 10 min. Chromogenic substrate solution (100 μl, 2 mM) was subsequently added to each well and the microtitre plate was incubated for a further 6 min at 37°C. The enzymatic reaction was stopped by adding 50 μl of 25% acetic acid to each well, and the presence of LPS was determined by measuring absorbance at 405 nm in a Tecan Infinite 200 Pro multiwell plate reader. A standard curve was generated using an *E. coli* endotoxin standard stock solution, which enabled the conversion of A405nm values into concentrations of LPS.

### Determination of bactericidal activity of antibiotics

As described above, stationary-phase bacteria grown overnight were washed twice and added at a final inoculum of 10^8^ c.f.u. ml^-1^ to 3 ml CA-MHB containing colistin and/or murepavadin. Cultures were incubated with shaking (37°C, 180 r.p.m.) for up to 8 h. Bacterial survival was determined after 2, 4, 6, and 8 h by serially diluting cultures in 10-fold steps in 200 μl sterile PBS (VWR International, USA), before enumeration of c.f.u. counts on MHA plates.

### Murine lung infection model

The use of mice was performed under the authority of the UK Home Office outlined in the Animals (Scientific Procedures) Act 1986 after ethical review by Imperial College London Animal Welfare and Ethical Review Body (PPL 70/7969). Wild-type C57BL/6 mice were purchased from Charles River (UK). All mice were female and aged between six to eight weeks. Mice were housed with five per cage with Aspen chip 2 bedding and 12 h light/dark cycles at 20–22 °C. Mice were randomly assigned to experimental groups. Water was provided *ad libitum* and mice were fed RM1 (Special Diet Services). To establish colonisation of the lungs, mice were anesthetized and intranasally inoculated with 10^7^ c.f.u. of *P. aeruginosa* PA14 in 50 μL of PBS, as described previously [59,60]. Infection was allowed to establish for 5 h, before mice were again anaesthetized and treated via the intranasal route with 50 μL of PBS alone, or PBS containing colistin (5 mg kg^-1^), murepavadin (0.25 mg kg^-1^), or a combination of colistin and murepavadin for 3 h. To enumerate bacterial load in the lungs, mice were humanely sacrificed, their lungs removed and homogenized in PBS, and then plated onto *Pseudomonas* isolation agar (Thermo Fisher Scientific).

### Statistical analyses

Experiments were performed on at least three independent occasions, and the resulting data are presented as the arithmetic mean of these biological repeats, unless stated otherwise. Error bars, where shown, represent the standard deviation of the mean. For single comparisons, a two-tailed Student’s *t*-test was used to analyse the data. For multiple comparisons at a single time point or concentration, data were analysed using a one-way analysis of variance (ANOVA) or a Kruskal-Wallis test. Where data were obtained at several different time points or concentrations, a two-way ANOVA was used for statistical analyses. Appropriate post-hoc tests (Dunnett’s, Tukey’s, Sidak’s, Dunn’s) were carried out to correct for multiple comparisons, with details provided in the figure legends. Asterisks on graphs indicate significant differences between data, and the corresponding p-values are reported in the figure legend. All statistical analyses were performed using GraphPad Prism 7 software (GraphPad Software Inc., USA).

## Acknowledgements

Aishwarya Krishna, Laura Nolan, Alain Filloux, Ollie Fletcher (all Imperial College London) and Alireza Abdolrasouli (Imperial College Healthcare NHS Trust) are gratefully acknowledged for providing bacterial strains. Lisa Haigh (Imperial College London) is thanked for processing and analysing mass spectrometry samples. Vladimir Pelicic (Imperial College London) is thanked for providing access to the Axio Imager.A2 Zeiss microscope. A.S. is supported by a PhD studentship funded by a Medical Research Council Doctoral Training Award to Imperial College London (MR/N014103/1). A.K. is funded by the DFG German Research Foundation (KL3191/1-1) and the European Union’s Horizon 2020 research and innovation programme under the Marie Skłodowska-Curie Actions grant agreement “BacDrug” (838183). M.B. and M.M.S gratefully acknowledge support from the Rosetrees Trust (M300-CD1). D.A.I.M. and R.C.D.F. gratefully acknowledge funding from an MRC Career Development Award (MR/M009505/1). J.C.D is supported by the NIHR through a Senior Investigator award and the Imperial Biomedical Research Centre (BRC). R.M. & J.C.D. are supported by Cystic Fibrosis Trust funding. G.J.L-M. is funded by the MRC Confidence in Concept Fund and a ISSF Wellcome Trust Grant (105603/Z/14/Z). T.B.C. is a Sir Henry Dale Fellow jointly funded by the Wellcome Trust and Royal Society (107660/Z/15Z). L.E.E. and A.M.E. acknowledge funding from the Wellcome Trust (204337/Z/16/Z). A.M.E. also acknowledges support from the National Institute for Health Research (NIHR) Imperial Biomedical Research Centre (BRC), and declares funding from Shionogi & Co., Ltd for an unrelated project.

## Author Contributions

A.S., T.B.C. and A.M.E. designed the experiments. A.S., K.L.H.H., A.K., M.B., L.E.E., R.C.D.F., G.J.L-M., D.A.I.M. and T.B.C. performed experiments or generated bacterial strains or reagents. All authors analysed data and contributed to writing of the manuscript.

