## Supplementary information for "Colistin kills bacteria by targeting lipopolysaccharide in the cytoplasmic membrane"

**This file includes:**

Supplementary Tables 1-2

Supplementary Figures 1-11

| Bacterial Strain | Description | Source/Reference |
|---|---|---|
| *E. coli* | | |
| MC1000 pEmpty | Background strain (*araD139*, Δ(ara, leu)7697, Δ*lacX74*, *galU, galK, strA*) harbouring the IPTG-inducible pDM1 plasmid (GenBank MN128719) | Dortet *et al.*, 2018 [34] |
| MC1000 *mcr-1* | MC1000 strain harbouring the pDM1 plasmid encoding the *mcr-1* gene amplified from a clinical *E. coli* isolate | Dortet *et al.*, 2018 [34] |
| *P. aeruginosa* | | |
| PA14 | Wild-type reference strain; highly virulent human isolate representing most common clonal group worldwide | Lee *et al.*, 2006 [61] |
| AK3 | Multi-drug resistant human clinical isolate from sputum of cystic fibrosis patient – mucoid strain | This study |
| AK10 | Multi-drug resistant human clinical isolate from sputum of cystic fibrosis patient | This study |
| AK20 | Multi-drug resistant human clinical isolate from sputum of cystic fibrosis patient | This study |
| AK6 | Multi-drug resistant human clinical isolate from sputum of cystic fibrosis patient – mucoid strain | This study |
| AK12 | Multi-drug resistant human clinical isolate from sputum of cystic fibrosis patient – mucoid strain | This study |
| AK8 | Multi-drug resistant human clinical isolate from sputum of cystic fibrosis patient – mucoid strain | This study |
| AK17 | Multi-drug resistant human clinical isolate from sputum of cystic fibrosis patient | This study |
| AK9 | Multi-drug resistant human clinical isolate from sputum of cystic fibrosis patient | This study |
| AK14 | Multi-drug resistant human clinical isolate from sputum of cystic fibrosis patient | This study |
| AK5 | Multi-drug resistant human clinical isolate from sputum of cystic fibrosis patient – mucoid strain | This study |
| AK11 | Multi-drug resistant human clinical isolate from sputum of cystic fibrosis patient – mucoid strain | This study |
| AK13 | Multi-drug resistant human clinical isolate from sputum of cystic fibrosis patient | This study |
| AK18 | Multi-drug resistant human clinical isolate from sputum of cystic fibrosis patient | This study |
| AK22 | Multi-drug resistant human clinical isolate from sputum of cystic fibrosis patient | This study |
| AK19 | Multi-drug resistant human clinical isolate from sputum of cystic fibrosis patient – murepavadin-resistant mucoid strain | This study |
| AK21 | Multi-drug resistant human clinical isolate from sputum of cystic fibrosis patient – murepavadin-resistant | This study |

**Supplementary Table 1. Bacterial strains used in this study.**

| Bacterial Strain | Colistin MIC (µg/ml) | Murepavadin MIC (µg/ml) |
|---|---|---|
| PA14 | 0.5 | 0.2 |
| AK3 | 1 | 0.8 |
| AK10 | 1 | 1.6 |
| AK20 | 1 | 1.6 |
| AK6 | 1 | 1.6 |
| AK12 | 2 | 0.8 |
| AK8 | 2 | 1.6 |
| AK17 | 1 | 1.6 |
| AK9 | 1 | 1.6 |
| AK14 | 2 | 0.8 |
| AK5 | 2 | 1.6 |
| AK11 | 1 | 1.6 |
| AK13 | 1 | 1.6 |
| AK18 | 1 | 1.6 |
| AK22 | 1 | 0.8 |
| AK19 | >8 | >3.2 |
| AK21 | 4 | >3.2 |

**Supplementary Table 2. Minimum inhibitory concentrations (MICs) of colistin and murepavadin against the *P. aeruginosa* strains used in this study.** MICs were determined in cation-adjusted Mueller-Hinton Broth (CA-MHB) using the broth microdilution method, as previously described [56]. The MIC was defined as the lowest antibiotic concentration at which there was no visible growth of bacteria after 18 hours incubation. Data are presented as the median of at least 3 independent biological replicates.

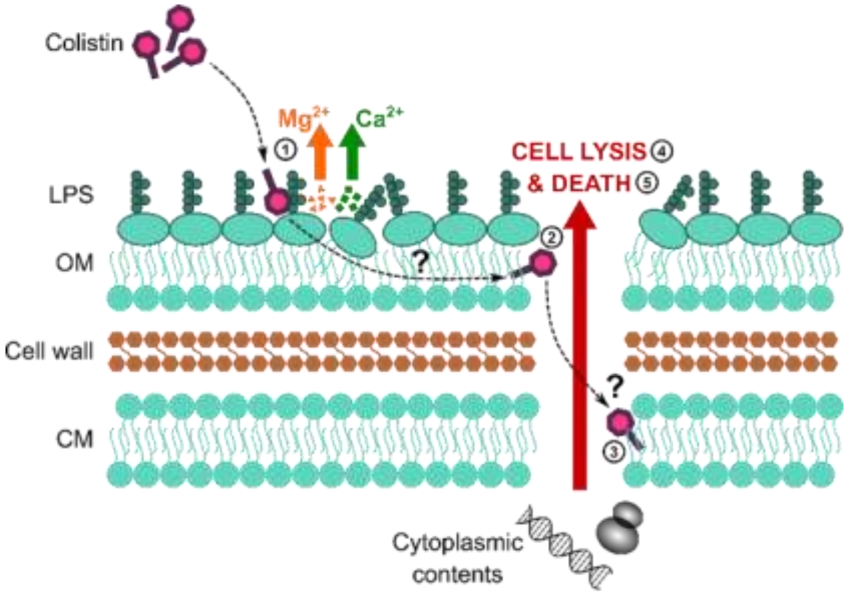


**Supplementary Figure 1. Colistin causes outer membrane (OM) disruption, but the process by which this leads to cytoplasmic membrane (CM) damage and bacterial lysis is not known.** Diagrammatic representation of the current hypothesised mechanism of action of colistin: (1) Colistin binds to LPS in the OM, causing displacement of cations that form bridges between LPS molecules and leading to membrane disruption. (2) The antibiotic then crosses the OM via a process termed 'self-directed uptake'. (3) Colistin subsequently disrupts the CM through a mechanism that remains unclear. (4,5) Once the cell envelope is permeabilized, the bacteria lyse and are killed.

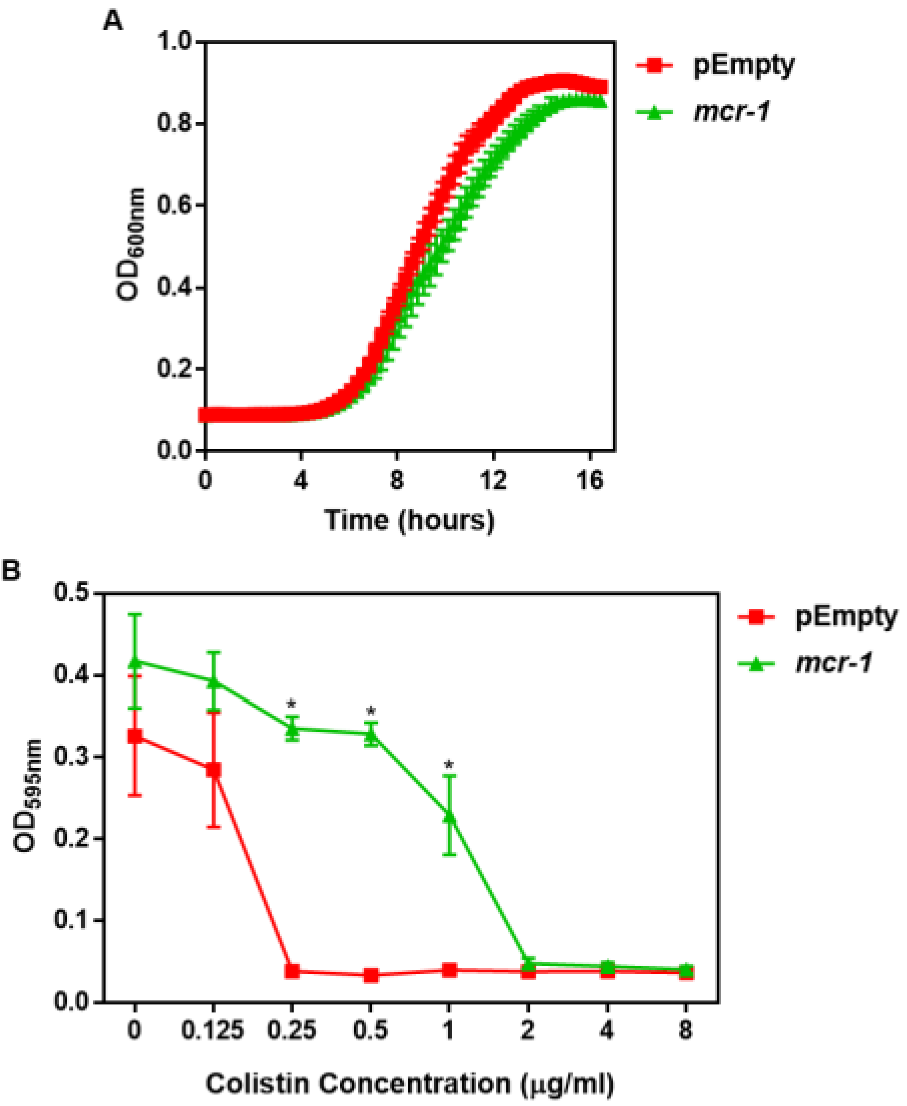


**Supplementary Figure 2. Characterisation of the *E. coli* MC1000 strain harbouring a plasmid encoding the colistin resistance gene *mcr-1*, and an MC1000 strain containing the pDM1 plasmid only (pEmpty) as a control strain. A,** Growth of *E. coli* MC1000 cells containing an empty pDM1 plasmid (pEmpty) or MC1000 cells containing the pDM1 plasmid expressing the *mcr-1* gene, as determined by measuring $OD_{600nm}$ over 16 hours incubation at 37°C (n=3 in triplicate). **B,** Final growth densities of *E. coli* MC1000 cells with an empty pDM1 plasmid and MC1000 cells harbouring the pDM1 plasmid encoding the *mcr-1* gene in CA-MHB media containing the indicated concentrations of colistin, as determined by measuring $OD_{595nm}$ after 18 hours incubation (n=3 in triplicate; *p<0.0001 compared to pEmpty strain). Expression of the mobilised colistin resistance determinant *mcr-1* by *E. coli* MC1000 bacteria resulted in only a minor growth defect relative to empty plasmid-containing control cells **(A)**. As expected, *E. coli* MC1000 cells producing MCR-1 had an 8-fold increase in the MIC of colistin in comparison to the isogenic empty plasmid strain (2 µg $ml^{-1}$ versus 0.25 µg $ml^{-1}$) **(B)**. This confirmed that the bacteria harbouring a plasmid with the *mcr-1* gene were resistant to colistin. Data in **B** were analysed by a two-way ANOVA with Sidak's post-hoc test. Data are presented as the arithmetic mean, and error bars represent the standard deviation of the mean.

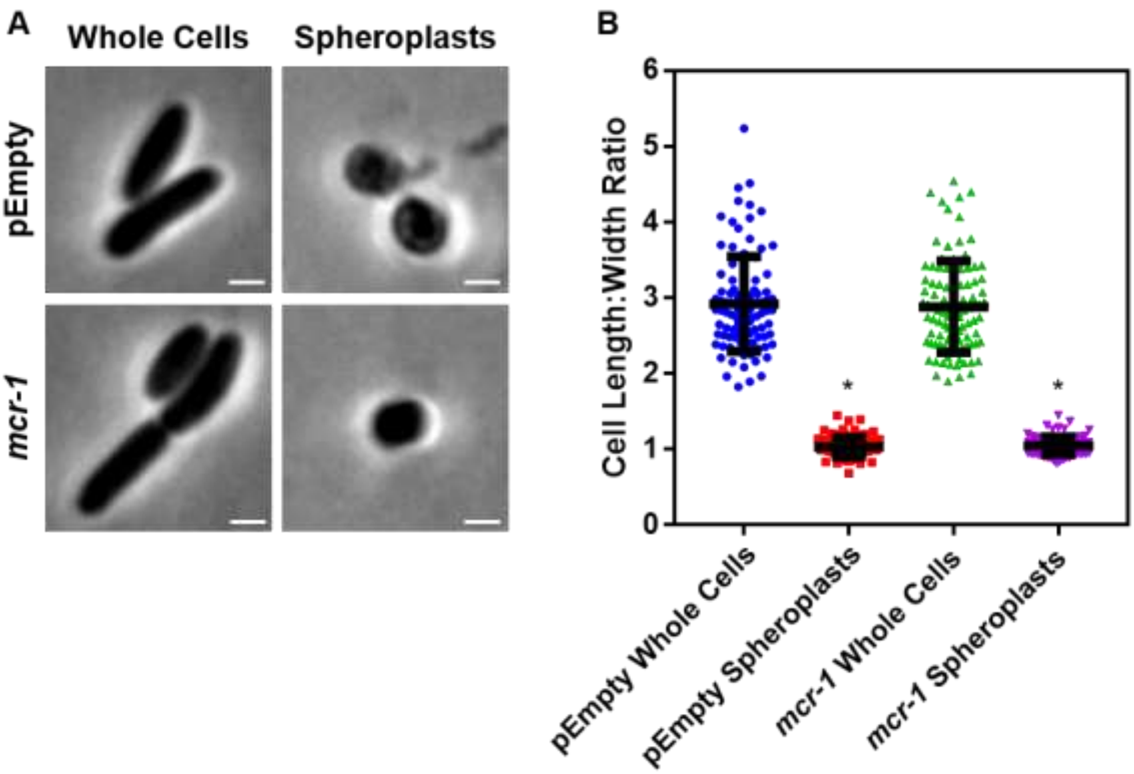


**Supplementary Figure 3. Formation of *E. coli* pEmpty and *mcr-1* spheroplasts. A,** Representative phase contrast micrographs of *E. coli* MC1000 cells harbouring an empty pDM1 plasmid (pEmpty) or a pDM1 plasmid expressing the colistin-resistance determinant *mcr-1* before (*Whole Cells*) and after (*Spheroplasts*) treatment with 0.25 mg $ml^{-1}$ EDTA (to remove the outer membrane) and 1 mg $ml^{-1}$ lysozyme (to remove the cell wall) for 1 hour at 30°C, followed by the addition of trypsin (0.5 mg $ml^{-1}$) for 15 minutes, with all incubations occurring in Tris buffer (0.03 M, pH 8.0) containing 20% sucrose (Scale bars: 5 µm). **B,** Quantification of length:width ratio of *E. coli* MC1000 cells containing an empty plasmid, or *E. coli* cells with the same plasmid encoding the *mcr-1* gene, before (*Whole Cells*) and after (*Spheroplasts*) exposure to EDTA, lysozyme and trypsin **(A)** (n=100 cells per group; *$p<0.0001$ compared to the respective Whole Cells). Removal of the outer membrane and cell wall from both *E. coli* MC1000 pEmpty and *mcr-1*-expressing bacteria results in the formation of round cells with a length:width ratio of ~1 **(A, B)**, confirming the successful production of *E. coli* spheroplasts lacking the outer membrane and cell wall. Data in **B** were analysed by a one-way ANOVA with Tukey's post-hoc test. Data are presented as the arithmetic mean, and error bars represent the standard deviation of the mean.

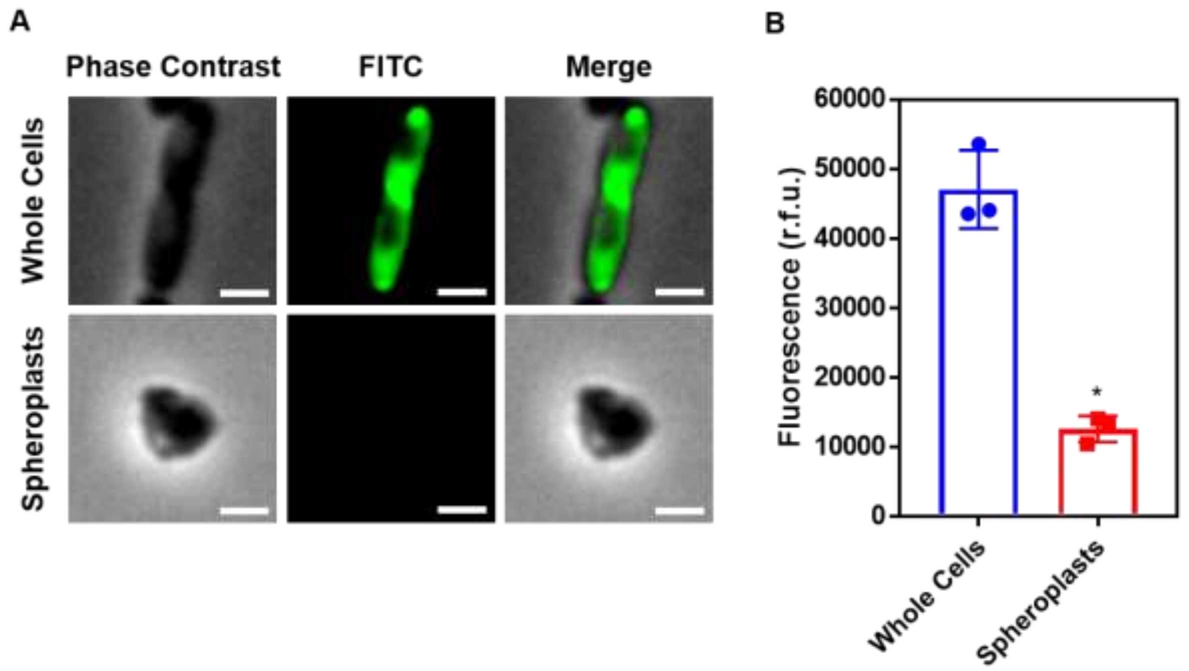


**Supplementary Figure 4. Conversion of *E. coli* whole cells to spheroplasts results in removal of the OM, and no OM contamination in the CM. A,** Representative fluorescence microscopy images of *E. coli* MC1000 pEmpty cells labelled with fluorescein isothiocyanate (FITC, 0.5 mg $ml^{-1}$) before (*Whole Cells*) and after (*Spheroplasts*) conversion to spheroplasts with EDTA (0.25 mg $ml^{-1}$) and lysozyme (1 mg $ml^{-1}$) in Tris buffer (0.03 M, pH 8.0) containing 20% sucrose, as described in Supplementary Fig. 3 (Scale bars: 5 μm). **B,** Quantification of fluorescence from FITC-labelled *E. coli* MC1000 pEmpty cells before (*Whole Cells*) and after (*Spheroplasts*) conversion to spheroplasts (n=3 in triplicate; *p<0.01 compared to Whole Cells). Proteins in the OM of whole *E. coli* MC1000 cells were tagged with a FITC fluorophore for 30 minutes, as previously described [57]. Following conversion of these labelled bacterial cells to spheroplasts, there was virtually no fluorescence from FITC visible by microscopy **(A)**, or when quantifying the entire cell population **(B)**. This confirmed that the OM had been successfully removed during the formation of spheroplasts, and that there was no contamination of the CM with material from the OM. Data in **B** were analysed by a paired Student's *t*-test. Data are presented as the arithmetic mean, and error bars represent the standard deviation of the mean.

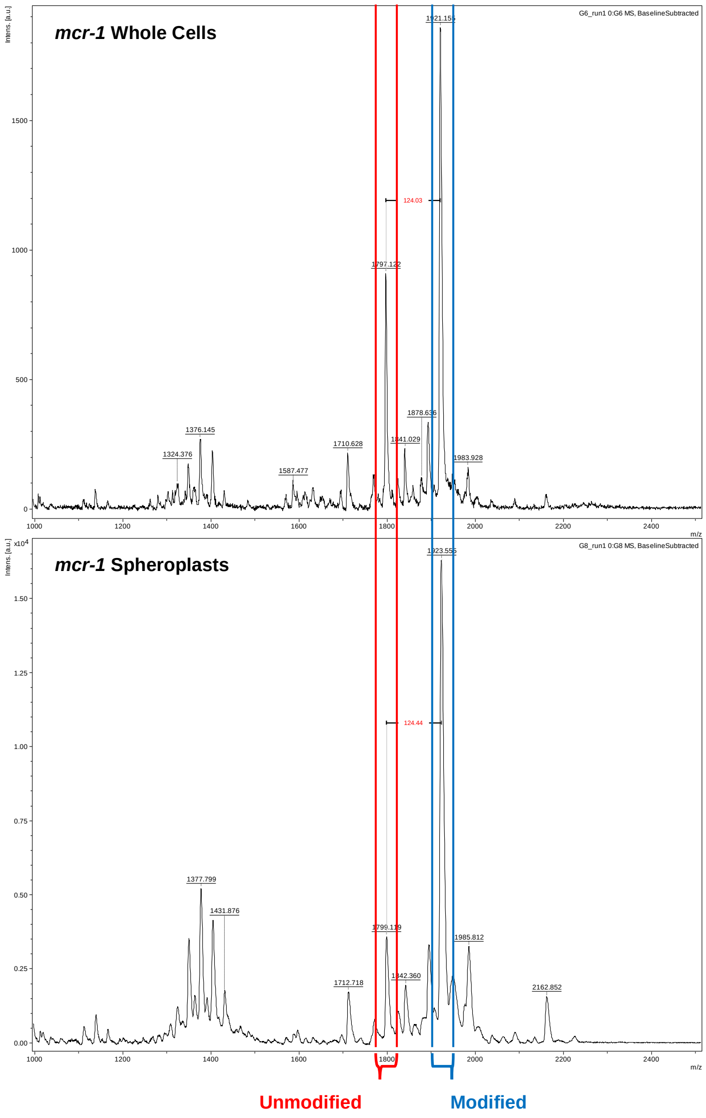


**Supplementary Figure 5. The ratio of modified lipid A to unmodified lipid A is significantly greater in the cytoplasmic membrane than in the outer membrane of *E. coli* expressing *mcr-1*.** Representative mass spectra showing the ratio of unmodified lipid A (red) to lipid A modified with phosphoethanolamine (blue) in whole cells and spheroplasts of *E. coli* MC1000 expressing *mcr-1*, as determined by MALDI-TOF-based lipidomics. There was a higher proportion of LPS modified with pETN in *mcr-1*-expressing *E. coli* MC1000 spheroplasts compared to whole *E. coli* MC1000 cells, demonstrating that there is an increased abundance of unmodified LPS in the OM compared to the CM.

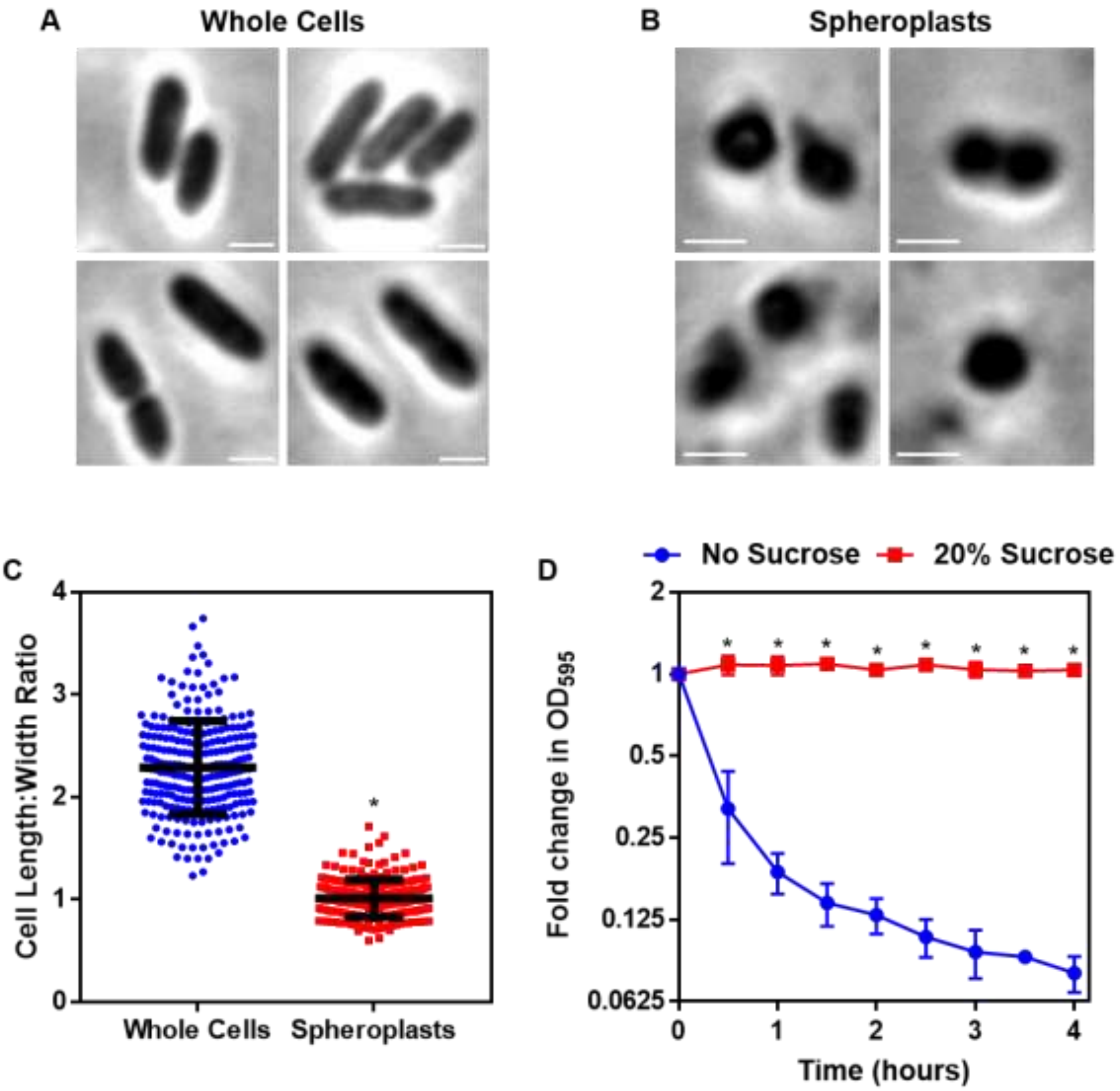


**Supplementary Figure 6. Formation of *P. aeruginosa* spheroplasts. A, B,** Representative phase contrast micrographs of *P. aeruginosa* PA14 cells before **(A)** and after **(B)** treatment with 0.25 mg $ml^{-1}$ EDTA (to remove the OM) and 1 mg $ml^{-1}$ lysozyme (to remove the cell wall) for 1 hour at 30°C, followed by the addition of trypsin (0.5 mg $ml^{-1}$) for 15 minutes, with all incubations occurring in Tris buffer (0.03 M, pH 8.0) containing 20% sucrose (Scale bars: 5 µm). **C,** Quantification of length:width ratio of *P. aeruginosa* PA14 whole cells **(A)** and *P. aeruginosa* spheroplasts following exposure to EDTA, lysozyme and trypsin **(B)** (n=250 cells per group; *p<0.0001 compared to Whole Cells). **D,** Lysis of *P. aeruginosa* spheroplasts exposed to EDTA (0.25 mg $ml^{-1}$) and lysozyme (1 mg $ml^{-1}$) at 30°C in Tris buffer (0.03 M, pH 8.0), in the presence or absence of 20% sucrose, as determined by $OD_{595nm}$ measurements (n=4; *p<0.0001 compared to No Sucrose). After removal of the OM and cell wall with EDTA and lysozyme respectively, the resulting *P. aeruginosa* cells formed had a length:width ratio of almost exactly 1 **(C)** and immediately lysed when not maintained in an osmoprotective environment containing 20% sucrose **(D)**. Together, these data confirmed the successful production of *P. aeruginosa* spheroplasts lacking the OM and cell wall. Data in **C** were analysed by an unpaired Student's *t*-test. Data in **D** were analysed by a two-way ANOVA with Sidak's post-hoc test. Data are presented as the arithmetic mean, and error bars represent the standard deviation of the mean.

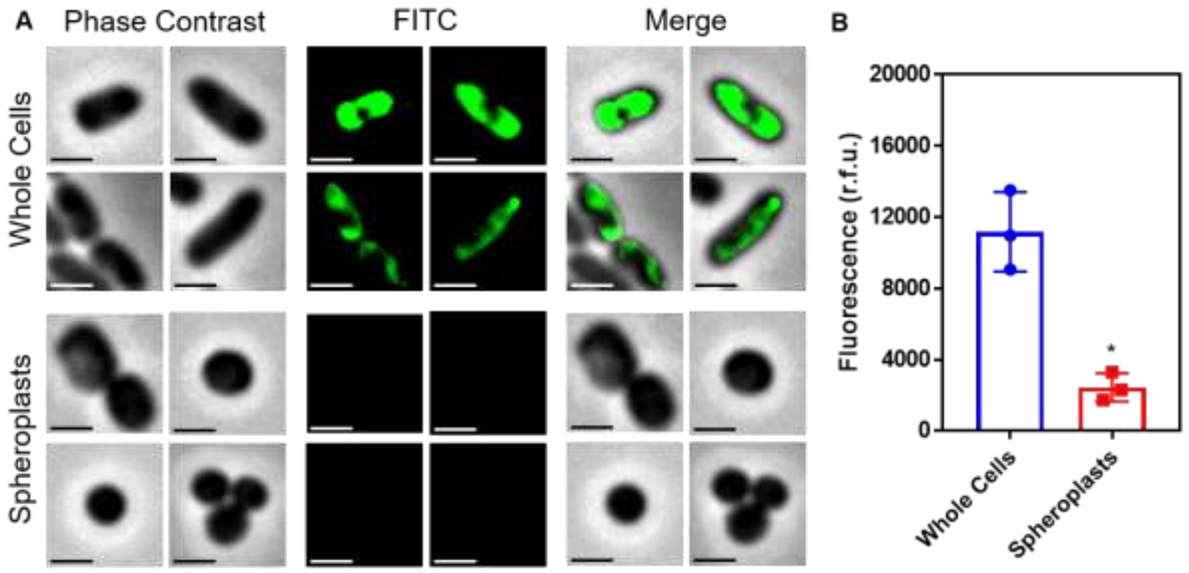


**Supplementary Figure 7. Conversion of *P. aeruginosa* whole cells to spheroplasts results in removal of the OM, and no OM contamination in the CM. A,** Representative fluorescence microscopy images of *P. aeruginosa* PA14 cells labelled with fluorescein isothiocyanate (FITC, 0.5 mg $ml^{-1}$) before (*Whole Cells*) and after (*Spheroplasts*) conversion to spheroplasts with EDTA (0.25 mg $ml^{-1}$) and lysozyme (1 mg $ml^{-1}$) in Tris buffer (0.03 M, pH 8.0) containing 20% sucrose, as described in Supplementary Fig. 6 (Scale bars: 5 μm). **B,** Quantification of fluorescence from FITC-labelled *P. aeruginosa* PA14 cells before (*Whole Cells*) and after (*Spheroplasts*) conversion to spheroplasts (n=3 in triplicate; *$p<0.05$ compared to Whole Cells). Proteins in the OM of whole *P. aeruginosa* cells were tagged with a FITC fluorophore for 30 minutes, as previously described [57]. Following conversion of these labelled bacterial cells to spheroplasts, there was virtually no fluorescence from FITC visible by microscopy **(A)**, or when quantifying the entire cell population **(B)**. This confirmed that the OM had been successfully removed during the formation of spheroplasts, and that there was no contamination of the CM with material from the OM. Data in **B** were analysed by a paired Student's *t*-test. Data are presented as the arithmetic mean, and error bars represent the standard deviation of the mean.

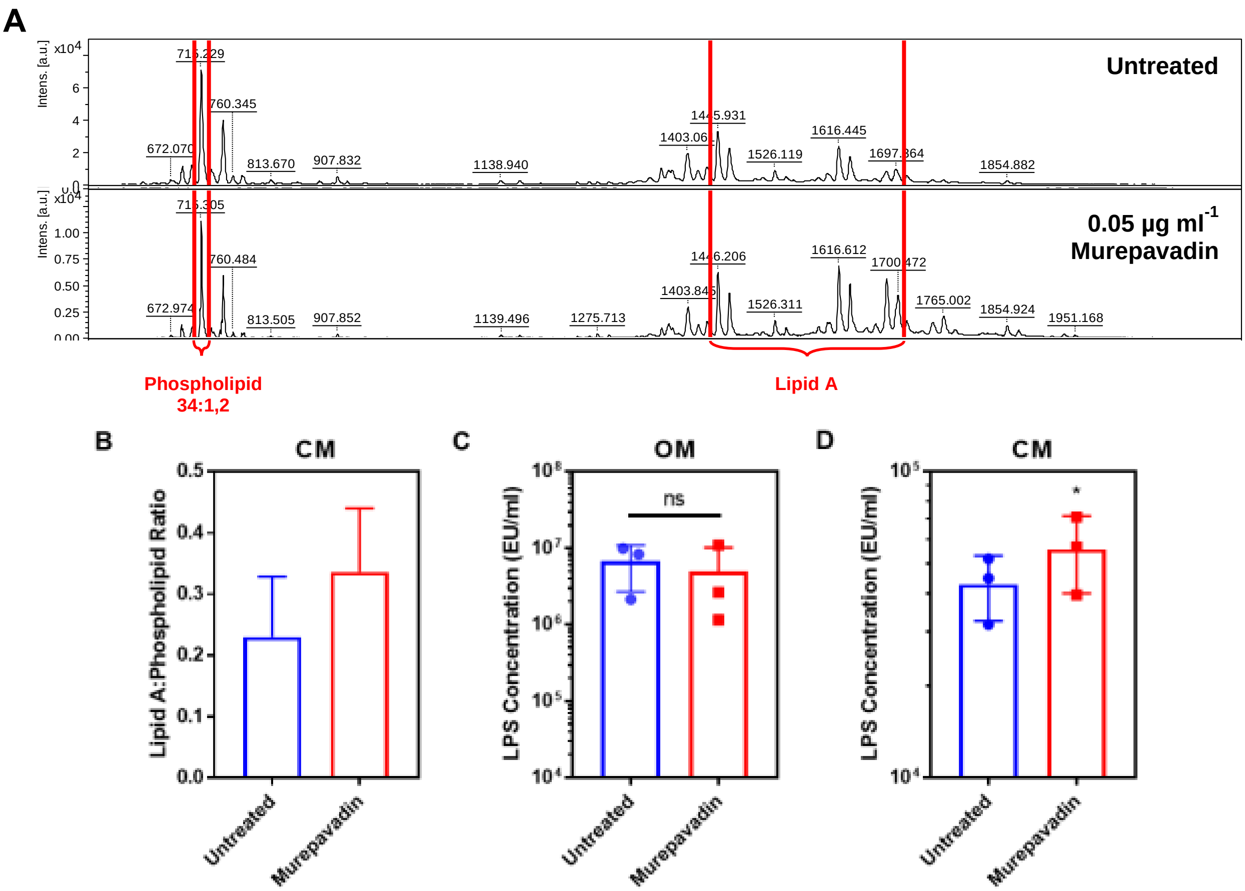


**Supplementary Figure 8. Murepavadin increases the abundance of LPS in the cytoplasmic membrane of *P. aeruginosa*. A, B,** Representative mass spectra **(A)** and quantification **(B)** showing the ratio of lipid A to a membrane phospholipid species (34:1,2) in the CM of *P. aeruginosa* PA14 spheroplasts exposed, or not, to murepavadin (0.05 µg $ml^{-1}$) for 2 hours, as determined by MALDI-TOF-based lipidomics (n=3 in duplicate). **C, D,** Quantification of LPS levels in the OM **(C)** and CM **(D)** of *P. aeruginosa* PA14 during conversion to spheroplasts from whole cells pre-exposed, or not, to the LPS transport inhibitor murepavadin (0.05 µg $ml^{-1}$) for 2 hours, as determined using the Limulus Amebocyte Lysate (LAL) assay (n=3; ns: $p>0.05$, *$p<0.05$ compared to Untreated conditions). Exposure of *P. aeruginosa* to a sub-lethal concentration of the LPS transport inhibitor murepavadin triggered an accumulation of LPS in the CM, but had no effect on LPS levels in the OM. Furthermore, MALDI-TOF analysis confirmed that the LPS in murepavadin-treated cells was unmodified, confirming it could be accurately detected and quantified using the LAL assay. Data in **B-D** were analysed by a two-tailed paired Student's *t*-test. Data are presented as the arithmetic mean, and error bars represent the standard deviation of the mean.

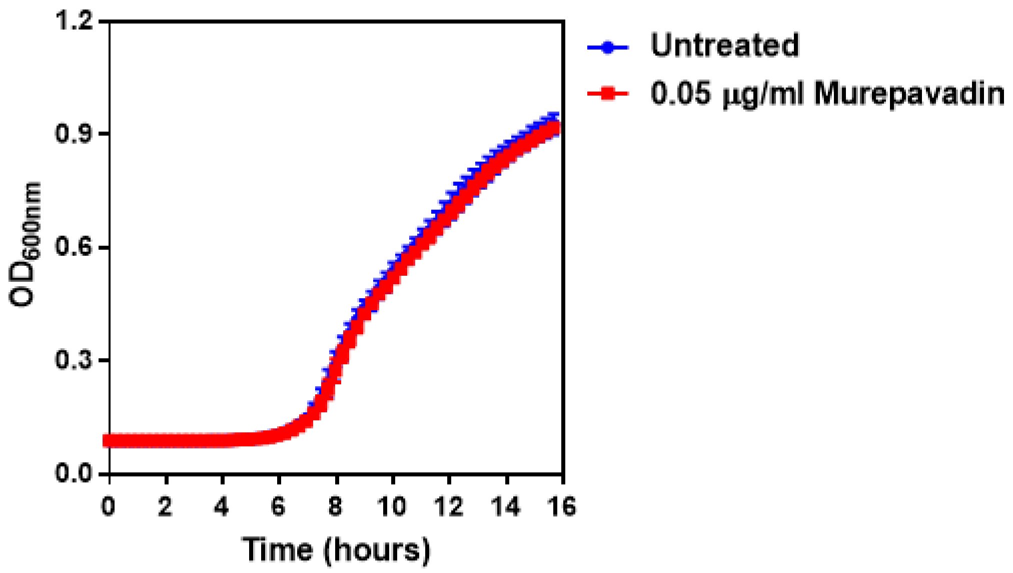


**Supplementary Figure 9. The LPS transport inhibitor murepavadin has no effect on reducing growth of *P. aeruginosa* at the concentration used.** Growth of *P. aeruginosa* PA14 cells in the presence of a sub-lethal concentration of the LptD inhibitor murepavadin at the indicated concentration, as determined by measuring $OD_{600nm}$ during 16 hours incubation at 37°C (n=4; murepavadin-treated population not significantly different i.e. $p>0.05$ compared to Untreated). Exposure of *P. aeruginosa* to murepavadin at the concentration used in experiments had no effect on blocking bacterial growth, confirming that this was a sub-lethal concentration. Data were analysed by a two-way ANOVA with Sidak's post-hoc test. Data are presented as the arithmetic mean, and error bars represent the standard deviation of the mean.

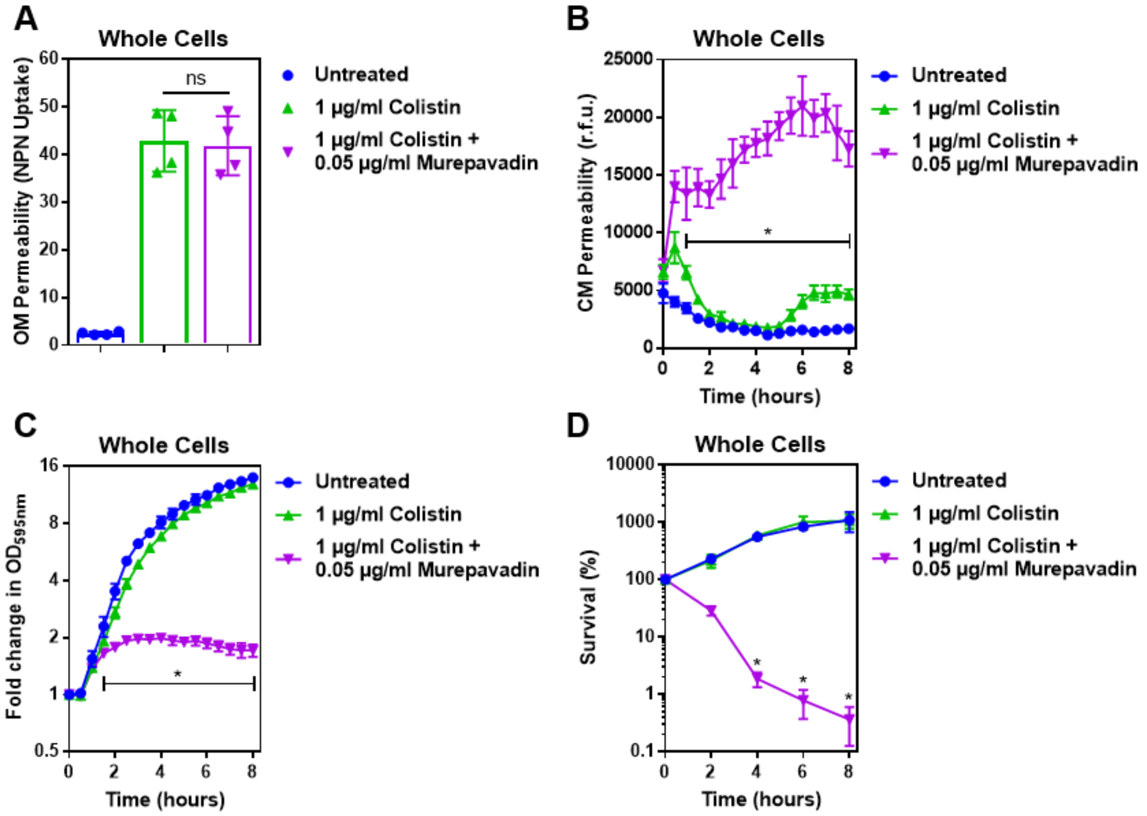


**Supplementary Figure 10. Murepavadin potentiates the activity of a sub-lethal colistin concentration, leading to enhanced CM damage, cell lysis and bacterial killing. A,** OM disruption of *P. aeruginosa* PA14 cells during 10 minutes exposure to colistin (1 µg $ml^{-1}$) in the absence or presence of murepavadin (0.05 µg $ml^{-1}$), as assessed by uptake of the fluorescent dye NPN (10 µM) (n=4, each data point represents the arithmetic mean of 20 replicate measurements; ns: p>0.05 between colistin-treated bacteria). **B,** CM disruption of *P. aeruginosa* PA14 cells exposed to colistin (1 µg $ml^{-1}$) in the absence or presence of murepavadin (0.05 µg $ml^{-1}$), as determined using 2.5 µM PI (n=4; *p<0.0001 for murepavadin-exposed cells compared to Colistin alone). **C,** Growth of *P. aeruginosa* PA14 cells exposed to colistin (1 µg $ml^{-1}$) in the absence or presence of murepavadin (0.05 µg $ml^{-1}$), as measured by $OD_{595nm}$ readings (n=4; *p<0.01 for murepavadin-exposed cells compared to Colistin alone). **D,** Survival of *P. aeruginosa* PA14 cells exposed to colistin (1 µg $ml^{-1}$) in the absence or presence of murepavadin (0.05 µg $ml^{-1}$), as determined by c.f.u. counts (n=4; *p<0.0001 for murepavadin-exposed cells compared to Colistin alone). Exposure of *P. aeruginosa* to murepavadin, which leads to accumulation of LPS in the CM, had no impact on the ability of colistin to damage the OM, further confirming that the LPS transport inhibitor, at the concentration used, did not significantly affect LPS levels in the OM **(A)**. However, a normally sub-lethal concentration of colistin was able to permeabilize the CM of murepavadin-exposed whole cells of *P. aeruginosa* **(B)**. This enhanced CM disruption by a sub-lethal concentration of colistin in combination with murepavadin led to an increase in lysis of *P. aeruginosa* cells, and ultimately greater bacterial killing **(C, D)**. Together, these data showed that accumulation of LPS in the CM via murepavadin enhanced the bactericidal activity of colistin, even at concentrations of the polymyxin antibiotic that otherwise had no growth inhibitory effects. Data in **A** were analysed by a one-way ANOVA with Tukey's post-hoc test. Data in **B, C** were analysed by a two-way ANOVA with Dunnett's post-hoc test. Data in **D** were analysed by a two-way ANOVA with Tukey's post-hoc test. Data are presented as the arithmetic mean, and error bars represent the standard deviation of the mean.

| Strain | Amikacin | Azithromycin | Aztreonam | Ceftazidime | Chloramphenicol | Ciprofloxacin | Colistin | Cotrimoxazole | Fosfomycin | Gentamicin | Levofloxacin | Meropenem | Minocycline | Pip/Tazobactam | Temocillin | Tetracycline | Timentin | Tigecycline | Tobramycin |
|---|---|---|---|---|---|---|---|---|---|---|---|---|---|---|---|---|---|---|---|
| AK3 | S | | I | S | R | R | S | R | | S | | S | | S | | | | | S |
| AK10 | S | | I | R | R | S | S | R | | S | | R | | S | | | | | S |
| AK20 | S | | I | S | S | S | S | S | | S | | S | | S | | | | | S |
| AK6 | R | | R | | | | S | R | R | R | R | R | S | S | S | | | | R |
| AK8 | S | | R | S | R | S | S | R | | S | | S | | S | | | | | S |
| AK12 | S | | I | S | R | S | S | R | | S | | S | | S | | | | | S |
| AK9 | R | | S | S | R | R | S | R | | R | | S | | S | | | | | R |
| AK14 | S | | S | S | R | S | S | R | | S | | S | | S | | | | | S |
| AK17 | S | | I | S | | S | S | S | | S | | S | | S | | | | | S |
| AK5 | S | | I | S | R | R | S | R | | R | | S | | S | | | | | S |
| AK11 | S | | S | S | S | S | S | S | | S | | S | | S | | | | | S |
| AK13 | S | R | R | R | R | R | S | R | S | R | | R | R | R | R | | | R | |
| AK18 | S | | R | R | R | R | S | R | S | R | R | R | S | S | R | | | | S |
| AK22 | S | | S | S | R | S | S | R | | S | | S | | S | | | | | S |
| AK19 | R | R | R | R | S | R | R | S | R | R | | R | S | R | R | S | | S | R |
| AK21 | R | R | R | R | S | R | R | S | R | R | | S | S | S | R | R | S | S | R |

**Supplementary Figure 11. Antibiogram summarising the antimicrobial susceptibilities of a panel of MDR *P. aeruginosa* human clinical strains isolated from sputum samples of cystic fibrosis patients.** S: susceptible (green), I: intermediate (orange), R: resistant (red). Bacterial strains were isolated and antimicrobial susceptibility testing was performed at the Royal Brompton Hospital, Royal Brompton and Harefield NHS Foundation Trust.
